# IRF5 regulates microglial myelin clearance and cholesterol metabolism after demyelination

**DOI:** 10.1101/2023.08.14.553274

**Authors:** Alejandro Montilla, Alazne Zabala, Ibai Calvo, Paloma Mata, Irene Tomé, Mirjam Koster, Amanda Sierra, Susanne M. Kooistra, Federico Nicolás Soria, Bart J.L. Eggen, Olatz Fresnedo, José Andrés Fernández, Vanja Tepavcevic, Carlos Matute, María Domercq

## Abstract

Interferon regulatory factor 5 (IRF5), a transcription factor highly involved in innate immunity that drives microglia/macrophage towards a pro-inflammatory state, has been associated to multiple sclerosis susceptibility but its role in MS pathogenesis is unknown. Here we analysed the role of IRF5 in multiple sclerosis animal models. *Irf*5^-/-^ mice showed exacerbated damage in the chronic phase of experimental autoimmune encephalomyelitis (EAE) mice, despite an initial delay in its onset, as well as after lysolecithin injection into the spinal cord. Transcriptomic and lipidomic analysis evidence a role of this transcription factor in myelin metabolism and cholesterol homeostasis. Indeed, *Irf*5^-/-^ mice showed an aberrant accumulation of myelin debris and lipidic structures, such as CE-containing lipid droplets and cholesterol crystals, suggesting that myelin-derived lipids are not properly processed. Cholesterol crystal accumulation leads to an aberrant inflammatory response, which block oligodendrocyte migration into the core of demyelinated lesion and remyelination. Pharmacologically facilitating cholesterol transport reduces lipid droplet accumulation and ameliorates EAE exacerbated damage in *Irf*5^-/-^ mice. These results reveal for the first time the role of Irf5, a transcription factor necessary to orchestrate the immune responses, in phagocytes lipid metabolism which could be pivotal in regenerative responses such as remyelination.

## INTRODUCTION

Multiple sclerosis (MS) is a chronic, inflammatory disease of the central nervous system (CNS) that involves demyelination and axonal degeneration, currently representing the most common cause of non-traumatic disability in young adults. Demyelinating focal plaques are the MS pathological hallmark, being mainly caused by immune cell infiltration across the blood-brain barrier (BBB) ^1, 2^. Nevertheless, demyelinating lesions can undergo myelin regeneration, in a process denominated remyelination. This process can partially reinstate saltatory conduction and resolve functional deficits, and is more efficient during the initial phases of the disease ^3^. Actually, a failure in the repairing mechanisms determines MS progression ^1^.

While microglial cells and infiltrating macrophages are key contributing to the demyelinating damages at MS early stages, they can also counteract these pathological processes by promoting remyelination. Indeed, efficient myelin clearance by specialized phagocytes is essential to facilitate regenerative processes ^4^; as free myelin inhibits oligodendrocyte progenitor cell (OPC) differentiation ^5, 6^, myelin debris need to be properly removed from the extracellular space not to interfere with the intrinsic mechanisms of repair. Nevertheless, not only myelin phagocytosis is crucial for remyelination, but also its proper intracellular processing ^7^. Microglia can redirect captured myelin to synthetize sterols, specific species that favour inflammation resolving by activating liver X receptor (LXR) ^8, 9^. Moreover, inefficient myelin degradation by aged phagocytes leads to the intracellular accumulation of cholesterol crystals, that in turn hinders regenerative mechanisms ^10^.

Microglial responses are dependent on the activation of different signaling pathways by pattern recognition receptors such as toll-like receptors (TLR). Indeed, TLR-associated intracellular mechanisms are essential for proper remyelination ^11^. IRF5 is a transcription factor that plays an important role in regulating immune responses downstream of TLRs, specifically, TLR-3, TLR-4, TLR-7, and TLR-9, and other innate immune receptors ^12, 13^. IRF5 plays a major role in inflammation, in the induction of pro-inflammatory cytokines and chemokines and in defining the macrophage inflammatory phenotype ^13–15^. IRF5 is expressed in B cells, dendritic cells and monocytes/macrophages and in the CNS, its expression is mainly reduced to microglia cells ^16, 17^. Interestingly, *Irf5* gene has been proposed as a risk factor for several autoimmune disorders ^18, 19^, and different polymorphisms in this gene reached significant association to MS development in independent cohorts ^20, 21^. Moreover, IRF5 is involved in the transcriptional axis controlling microglial P2X4^+^ reactive state ^22^, which potentiates myelin phagocytosis and remyelination in the experimental autoimmune encephalomyelitis model (EAE) ^23^. However, its role in the pathology is still unknown. In this work, we analysed the role of IRF5 in the context of demyelination and its involvement in remyelination, inducing both EAE and lysolecithin-demyelinating lesions in *Irf5*^-/-^ mice. We described a novel role of IRF5 in microglial myelin phagocytosis and processing, that directly links to proper regenerative capacities after demyelinating damages.

## MATERIALS AND METHODS

### Human samples

Post-mortem optic nerve samples from 13 MS patients and 12 control subjects (who died from non-neurological diseases) were obtained under the management of the Netherlands Brain Bank. All patients and controls had previously given written approval for the use of their tissue, according to the guidelines of the Netherlands Brain Bank. The clinical characteristics of the different experimental groups have been previously described ^24^. For comparisons, MS samples were matched with control samples for age, sex, and post-mortem delay.

### Animals

All experiments were performed according to the procedures approved by the Ethics Committee of the University of the Basque Country (UPV/EHU). Animals were handled in accordance with the European Communities Council Directive. Animals were kept under conventional housing conditions (22 ± 2°C, 55 ± 10% humidity, 12-hour day/night cycle and with *ad libitum* access to food and water) at the University of the Basque Country animal unit. All possible efforts were made to minimize animal suffering and the number of animals used. Experiments included C57BL/6 wild-type (WT) mice and *Irf5*^-/-^ C57BL/6 mice, the latter kindly provided by Prof. Tak W. Mak from the Princess Margaret Cancer Centre, UHN (Toronto, Canada).

### EAE immunization

Different EAEs were induced in 8- to 10-week-old male or female WT and *Irf5*^-/-^ mice. Mice were immunized with 200 µg of myelin oligodendrocyte glycoprotein 35-55 (MOG35–55; MEVGWYRPFSRVVHLYRNGK) in incomplete Freund’s adjuvant (IFA; Sigma) supplemented with 8 mg/mL Mycobacterium tuberculosis H37Ra (Fisher). Pertussis toxin (500 ng; Sigma) was injected intraperitoneally on the day of immunization and 2 days later, to facilitate the development of the disease model. Motor symptoms were recorded daily and scored from 0 to 8, as described before ^25^.

After EAE, mice were euthanized and the tissues were dissected out and differentially processed in accordance to the subsequent experimental procedure. For immunohistochemistry, the lumbar region of the spinal cord, where lesions typically accumulate, was fixed by immersion for 4 hours in 4% paraformaldehyde (PFA) dissolved in 0.1 M phosphate buffer (PB, pH = 7.4), rinsed in phosphate-buffered saline (PBS) and then transferred to 15% sucrose in 0.1 M PB for at least 2 days for cryoprotection. Next, tissue was frozen in 15% sucrose - 7% gelatine solution in PBS, and cut in a Leica CM3050 S cryostat to obtained 12-µm coronal sections. For real-time quantitative polymerase chain reaction (qPCR), the cervical and thoracic regions of the spinal cord, as well as peripheral immune-related organs, such as spleen or lymph nodes, were flash frozen in dry ice. For flow cytometry, the whole spinal cord was isolated.

### Lysolecithin-induced demyelination

To analyze remyelination in *Irf5*^-/-^ mice, we performed LPC-induced demyelination in the spinal cord of both WT and knock-out male mice. The lesions were induced by stereotaxic injection of 0.5μL of 1% lysolecithin (LPC; Sigma) in saline solution, as previously described ^26, 27^. Briefly, animals were anesthetized by intraperitoneal injection of a solution of ketamine (100 mg/kg) and xylazine (10 mg/kg). The tissue covering the vertebral column was making two longitudinal incisions into the *longissimus dorsi*, and the intravertebral space of the 13^th^ thoracic vertebra was exposed by removing the connective tissue after fixing the animal in the stereotaxic frame. Dura mater was then pierced using a 30G needle, and LPC was injected via a Hamilton syringe attached to a glass micropipette using a stereotaxic micromanipulator.

The lesion specific site was marked with sterile charcoal so that the area of tissue at the center of the lesions could be unambiguously identified afterwards. Following LPC injection, the wound was sutured and mice were allowed to recover. Mice were euthanized 4 and 14 days after surgery, in order to assess the response to demyelination. After LPC-induced demyelination, mice were perfused with 2% PFA for 15-20 minutes and spinal cords were post-fixed in 2% PFA for another 30 minutes. Tissue was then processed the same way as EAE lumbar spinal cords.

### Immunohistochemistry

Coronal sections of spinal cords from control animals, EAE mice and mice with LPC demyelinating lesions were analyzed by immunohistochemistry (IHC). Primary antibodies used for immunofluorescence on these tissues include: mouse anti-myelin basic protein (MBP) (1:1000; #808401 BioLegend), rabbit anti-MBP (1:200; #AB980 Millipore), rabbit anti-Iba1 (1:500; #019-19741 Wako Chemicals), mouse anti-SMI32 (1:1000; #801701 BioLegend) rat anti-CD3 (1:50; #MCA1477 Bio-Rad), rat anti-CD45R (1:200; #557390 BD Bioscience), mouse anti-iNOS (1:100; # 610329 BD Biosciences), mouse anti-GFAP (1:40; #MAB3402 Millipore), mouse anti-Olig2 (1:1000; # MABN50 Millipore) and mouse anti-APC (1:200; # OP80 Millipore). These primary antibodies were subsequently detected by incubation with appropriate Alexa Fluor 488 or 594 conjugated goat antibodies (1:250; Invitrogen). Moreover, cell nuclei were stained using Hoechst 33258 (Sigma-Aldrich). For Oil red O staining of EAE and LPC-induced demyelinating lesions, manufacturer’s protocol was followed. Briefly, tissues were incubated with 60% isopropanol for 5 minutes followed by a staining step with 60% Oil Red O dissolved in isopropanol for 15 minutes. Excess of stain was then rinsed with distilled water and IHC was performed subsequently.

Images were acquired using a Leica TCS STED SP8 confocal microscope, a Zeiss LSM800 confocal microscope or a Pannoramic MIDI II slide scanner (3DHistech) with the same settings for all samples within one experimental group. For the visualization of cholesterol crystals, confocal reflection microscopy was performed on the tissues using a ZEISS LSM 800 Airyscan confocal microscope, as these structures strongly reflect the light of the excitation laser. All the image analysis was performed with the ImageJ software (National Institutes of Health; NIH). For histological analysis of EAE lesions, images of the whole section were obtained. Lesion extents, as well as axonal damage, were normalized to the total white matter (WM) area of each section. The lesion area was defined by the lack of MBP staining along with the accumulation of myelin debris (identified by an increase in MBP fluorescence), and the accumulation of Iba1^+^ cells was normalized to this lesioned extent. At least three sections were analyzed per animal. For phagocytosis analysis, we automatically detected myelin blobs using the Threshold tool with a Gaussian blur (radius=1) and a Yen threshold. Microglia were identified using the Threshold tool with Yen threshold, and the Analyze particles tool to fill in the phagocytic pouches. Next, we quantified phagocytosis as the % blobs within microglia. For the assessment of LPC-induced lesions, dorsal funiculus images were obtained using a 20x objective, and analyses were performed similarly to those in the EAE tissue. To evaluate the distribution of different cell types in relation to the demyelinating lesions, we analyzed the fluorescence intensity of the markers in radial profiles comprising both lesioned and non-lesioned WM. Fluorescence intensity was averaged between profiles and normalized to allow comparisons. Distance was normalized likewise. After manual ROI creation to delineate the lesions, profiles were automatically drawn with a fixed width (250 px) and length (1 diameter of lesion ROI).

### Quantitative RT-PCR

Total RNA from EAE lumbar spinal cords, spleens and lymph nodes was isolated using TRIzol (Invitrogen) following the manufacturer’s instructions. Afterwards, 2 µg of this RNA was used to perform a retrotranscription protocol, using SuperScript III Reverse Transcriptase (200 U/μL; Invitrogen) and random hexamers as primers (Promega).

qPCRs were conducted in a Bio-Rad Laboratories CFX96 real-time PCR detection system, using iTaq Universal SYBR Green Supermix (Bio-Rad), that includes SYBR Green as DNA-binding dye and iTaq DNA polymerase. The specific primers for different T cell subtypes were designed Primer Express software (Applied Biosystems) at exon junctions to avoid genomic DNA amplification. The cycling conditions comprised 3 min of polymerase activation at 95°C and 40 cycles consisting of 10 s at 95°C and 30 s at 60°C. The amount of cDNA was quantified using a standard curve from a pool of cDNA obtained from the different conditions of the experiment. Finally, the results were normalized using a normalization factor based on the geometric mean of housekeeping genes obtained for each condition using the geNorm v3.5 software ^28^.

### Bulk RNA sequencing

RNA-sequencing was performed on total microglial populations, isolated by fluorescence-activated cell sorting (FACS), from control spinal cords of WT and *Irf5*^-/-^ mice. In order to isolate the cells while maintaining their specific activation state, all sorting steps were performed at 4°C. Spinal cords were mechanically dissociated and nuclear cells were isolated from debris using a Percoll gradient. Single cell suspensions were incubated with TruStain FcX™ (anti-mouse CD16/32) antibodies for 15 minutes to block unspecific bindings, and then stained for 30 minutes with CD11b-FITC (1:200; #101206 BioLegend), CD45-PE (1:100; #103106 BioLegend), Ly6C-PE/Cy7 (1:300; #128017 BioLegend) and SYTOX AADvanced™ Ready Flow™ (Thermo Fisher), a viability marker. We identified microglial population as SYTOX^-^/CD11b^+^/CD45^low^/Ly6C^-^. Cells were collected in RNAprotect Cell Reagent (Qiagen), and total RNA was extracted using the RNeasy Plus Micro kit (Qiagen), following manufacturer’s instructions.

NEBNext Low Input RNA Library Prep Kit for Illumina was used to process samples (n = 4 and 3 for WT and *Irf5*^-/-^ microglia respectively) (GenomeScan, Leiden, The Netherlands). RNA concentration and quality was determined with a Fragment Analyzer. Next, cDNA was synthesized and amplified from poly-A tailed mRNA. Clustering and DNA sequencing using the NovaSeq6000 was performed according to manufacturer’s protocols, using a concentration of 1.1 nM DNA. At least 20 million paired-end reads were generated per sample, with a quality score of ≥ 30. Quality checks, reads trimming and alignment to the most recent mouse genome were also performed.

All downstream bioinformatic analyses were performed in RStudio (v2021.09.0). For the differential gene expression analysis, low and non-expressed genes were excluded. The Bioconductor package edgeR ^29^ (v3.34.1) was used for normalization using the timed mean of M-values (TMM) method, and for identification of the differentially expressed genes (DEGs) between the different experimental groups by fitting a generalized linear model. DEGs were identified as those with an adjusted p-value < 0.05 and a log (Fold Change) > 1. Gene ontology (GO) analysis of the recognized DEGs for every comparison was performed using the DAVID ^30^ and Metascape ^31^ web resources.

### MALDI-IMS

In order to evaluate the changes in the lipidomic signatures in the context of LPC-induced demyelination, 12 µm-thick coronal sections were obtained from WT and *Irf5*^-/-^ mice spinal cords at 14 dpi. Tissues were scanned using a MALDI-LTQ-Orbitrap XL (Thermo Fisher) in the Spectroscopy Unit of the University of the Basque Country (UPV/EHU), using the negative-ion mode for the m/z region where the most relevant lipid species appear (650-1200 Da). The sections were covered with a 1,5-diaminonaphtalene matrix ^32^, using an in-house designed sublimator, and introduced in the MALDI source. Data acquisition was performed with a spatial resolution of 100 µm/pixel and 60.000 at m/z = 400 mass resolution.

For the processing of the lipid signatures obtained, the obtained spectra were processed using a software developed in MatLab (MathWorks). Briefly, the peaks obtained were identified and filtered. Then, the different lipid signatures related to the diverse regions of the spinal cord were segmented using a k-means clustering method. The lipidomic profile of the LPC-induced lesion, as well as the peri-lesion area and the healthy white matter of 5 WT and 5 *Irf5*^-/-^ injected mice were extracted and subsequently compared.

### Primary microglia culture

Primary mixed glial cultures were prepared from the cerebral cortex of neonatal mice (P4-P6). After 10-15 days in culture, microglia were isolated by mechanical shaking (400 rpm, 1 h) and purified by plating them on non-coated bacterial grade Petri dishes (Thermo Fisher Scientific), as previously described ^33^. Microglial cells obtained through this method were cultured in DMEM (Gibco) supplemented with 10% FBS (Gibco), at different cellular densities in accordance with the following experimental procedure. These cultures were practically pure of microglial cells (>99%) ^34^.

### Myelin phagocytosis assay

Mouse myelin was isolated as previously described ^35^. Briefly, spinal cord was mechanically homogenized in 0.32 M sucrose and subjected to repeated sucrose gradient centrifugation and osmotic shocks to separate myelin from other cellular components. Myelin concentration was measured with Bradford assay and adjusted to 1 mg/mL. Then, myelin was labelled with Alexa488-NHS dye (A2000 Life Technologies) for 1 hour at RT in PBS (pH 8). Dyed myelin was dialyzed to remove dye excess and resuspended in PBS (pH 7.4).

For the assessment of microglial phagocytosis, myelin was vortexed for 60 seconds for fragmentation and added to microglia culture medium (1:200 dilution). To evaluate myelin endocytosis, WT and *Irf5*^-/-^ primary microglia were incubated with Alexa488-NHS-labeled myelin for 1 hour at 37°C, rinsed and immediately fixed with 4% PFA. To evaluate myelin degradation, microglial cells were incubated with this myelin for 1 hour, rinsed and fixed 24 hours later. Myelin was quantified on Iba1^+^ cells using ImageJ on individual microglial cells outlined with the Iba1 immunostaining as the defining parameter for the ROIs. At least 50 cells were analyzed from each experiment (*n* = 3 independent experiments performed in triplicates).

### Wound healing assay

In order to assess the migratory capacity of WT and *Irf5*^-/-^ microglia, cells were seeded in DMEM + 10% FBS in glass-bottom dishes (Ibidi), generating a confluent monolayer. The monolayer was scratched in a straight line using a sterile 200 mL pipette tip. To follow the migration of microglia towards the scratched area, we performed a 24-hour time-lapse of the cells using a BioStation IM-Q microscope (Nikon), maintaining the dishes at 37°C and 5% CO_2_ during the whole extent of the experiment. The percentage of the scratched area occupied by microglia was quantified in the initial image obtained, as well as in images after 12 and 24 hours.

### Lipid extraction and quantification

In order to assess lipid metabolism of WT and *Irf5*^-/-^ microglia, we challenged these primary cells with 25 μg/mL purified myelin for 48 hours. After treatment, excess of myelin was washed with PBS, and cells were subsequently scrapped and centrifuged. Cell pellets were resuspended in PBS and sonicated in 2 cycles of 10 seconds, with intervals of 10 seconds between cycles and 25% amplitude. For lipid extraction, a commonly used method was carried out ^36^. Briefly, 2 mL of chloroform and 4 mL of methanol were added to 40 µg of protein from the cell homogenates; this initial volume also included a mixture of lipid standards. Tubes were vigorously shaken for 2 minutes, and 2 mL of chloroform were then added to the mixture. After shaking for another minute, 3.2 mL of distilled water was added to the tubes, and another 1-minute vortex step was performed. This mixture was centrifuged at 1500 g for 10 minutes, at 4°C, to allow the separation of the aqueous and organic phases. The lower, organic phases containing the lipids were transferred to clean tubes. Lipids retained in the aqueous phase were re-extracted by adding a mixture of chloroform, methanol and distilled water and repeating the shaking and centrifuge steps, and the new organic phase was combined with the previously obtained one. Last, the solvent was evaporated using a Thermo Savant SC250 EXP SpeedVac vacuum concentrator to obtain the final lipid extract.

Lipids in this extract were analyzed using UltiMate 3000 ultrafast liquid chromatography system (UHPLC; Thermo Scientific) coupled to a QExactive™ HF-X Hybrid Quadrupole-Orbitrap mass spectrometer (MS), in the University of the Basque Country (UPV/EHU) facilities. The extract was resuspended in 90 µL of 9:1 methanol:toluene mixture, and 7 µL of the resulting supernatant was injected into the HPLC-MS system. Electrospray ionization was performed in either positive or negative ion mode. Lipid species predicted by the quantification of the HLPC-MS results were filtered, classified into families and quantified in accordance to the standard signals and their concentrations. Lipid ontology (LION) analysis of the differential lipids was performed using the LION/web application ^37^. This experiment was performed on *n* = 4 different cultures from WT and *Irf5*^-/-^ mice, and the data represent the mean quantity (µg) of each species in a lipid class, normalized to the quantity of initial protein from each sample.

### Statistical analysis

Data are presented as mean ± standard error of mean (SEM) and n represents the number of animals or cultures analyzed; this is specificized in figure legends. Statistical analyses were performed using GraphPad Prism 8 (GraphPad Software Inc), applying the corresponding statistical treatment for each experiment. Comparisons between two groups were analysed using paired Student’s two-tailed t-test for data coming from in vitro experiments, unpaired Student’s two-tailed test for data coming from in vivo experiments and Mann-Whitney U test in the case of comparisons regarding EAE neurological scores. Comparisons among multiple groups were analysed by one-way ANOVA followed by Bonferroni post-hoc analysis. In all instances, p values < 0.05 were considered as statistically significant.

## RESULTS

### Microglial *Irf5* expression is downregulated response to demyelination

Different *Irf5* polymorphisms have been linked to MS development in diverse independent cohorts ^20, 21^. Previous data in our laboratory showed that *Irf5*, as well as *P2x4*, were upregulated in total RNA from the spinal cord of MOG-immunized mice^20^. To address whether *Irf5* expression is altered during demyelination, we assessed its expression in tissues coming from MS patients. We did not detect differences in the expression of IRF5 transcription factor in total RNA from post-mortem optic nerve samples of control and MS patients ^25^, as analyzed by qPCR, (Fig. 1A), suggesting that this gene is not upregulated during the development or at late stages of the disease. However, an *in silico* analysis of available data obtained by single-cell RNA sequencing of tissues coming from healthy humans and active multiple sclerosis patients ^38^ showed that *Irf5* is downregulated in microglial cells in the pathology (Fig. 1B; *n* = 4). We further analyzed the global versus microglia specific expression of Irf5 after EAE immunization. Whereas an increase in Irf5 expression was detected in total spinal cord RNA both at EAE peak and chronic phase (postimmunization day 20 and 35 respectively; Fig. 1C), the specific expression of *Irf5* in FACS-isolated microglia (Cd11b^+^CD45^high^) decreased at all EAE stages (Fig. 1D).

**Figure 1.**
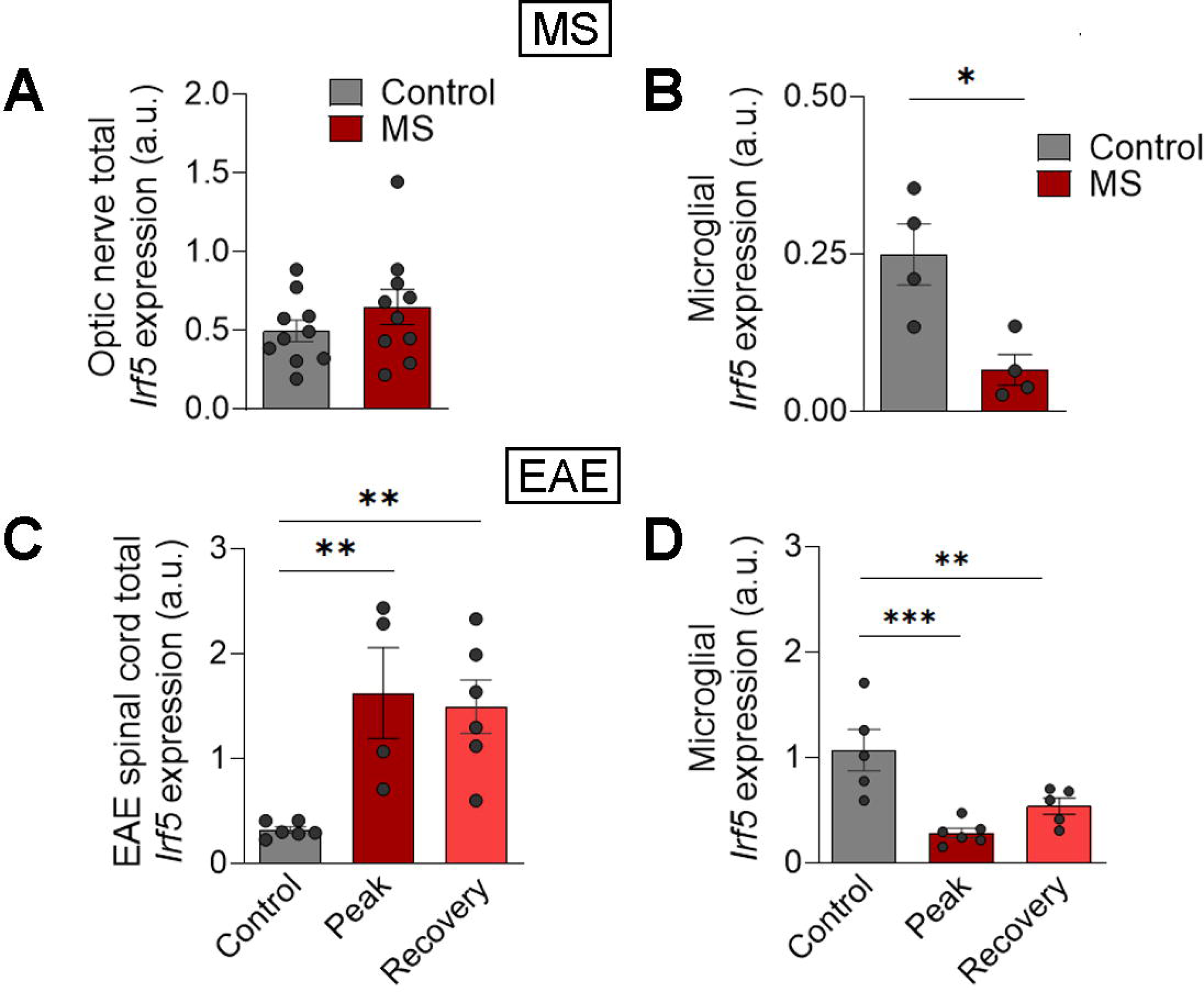
Microglial *Irf5* expression is decreased in MS patients and EAE mice. **(A)** Relative expression of *Irf5* in total mRNA isolated from post-mortem optic nerve samples of control and MS patients (*n* = 10). **(B)** *In silico* quantification of microglial *Irf5* expression in healthy and early active MS human brain tissues (*n* = 4). Raw data obtained from Gene Expression Omnibus (accession number: GSE124335). **(C)** Relative expression of *Irf5* in total mRNA isolated from spinal cord of control (n = 7) and EAE mice at the peak (n = 4) and recovery phases (*n* = 8). **(D)** Expression of *Irf5* in FACS isolated microglia (CD11b^+^/CD45^low^) from the spinal cord of control mice (n=4) and from EAE mice at chronic phase (*n* = 6). Data are presented as means ± SEM. *p < 0.05, **p < 0.005, ***p < 0.001.

These results highlight a specific microglial downregulation of *Irf5* in response to demyelination in human as well as in mice. In order to check its relevance in disease development and resolution, we induced different mouse models of demyelination and remyelination in *Irf5*^-/-^ mice.

### IRF5 deficiency exacerbates damage at EAE recovery phase

We first addressed the impact of *Irf5* deletion in EAE development. We observed a significant delay in the onset of motor symptoms in *Irf5*^-/-^ mice compared to WT ones (Fig. 2A, B), suggesting a role of this transcription factor in peripheral immune priming as previously stated for other IRF transcription factors ^39^. Despite this initial delay, *Irf5*^-/-^ mice showed no difference in the maximal neurological score at EAE peak (Fig. 2A-B). However, *Irf5*^-/-^ mice presented exacerbated neurological scores at EAE chronic phase (Fig. 2B) and a disruption of the regeneration capacities as the mice showed an increase in the time necessary to initiate recovery (Fig. 2B). At histological level, demyelinated lesions, defined by the presence of myelin loss or damage tends to be larger although not statistically significant (Fig. 2C; p = 0.074). However, we found a higher accumulation of Iba1^+^ cells inside the lesions and an increase in axonal damage (assessed with SMI32 marker) in *Irf5*^-/-^ mice (Fig. 2C). These observations confirm the exacerbation of tissue damage in *Irf5*^-/-^ mice at EAE chronic phase. Although the role of IRF5 in EAE priming would need further research, we concentrated on its impact in EAE chronic phase because the data point to a novel role of this factor in regeneration.

**Figure 2.**
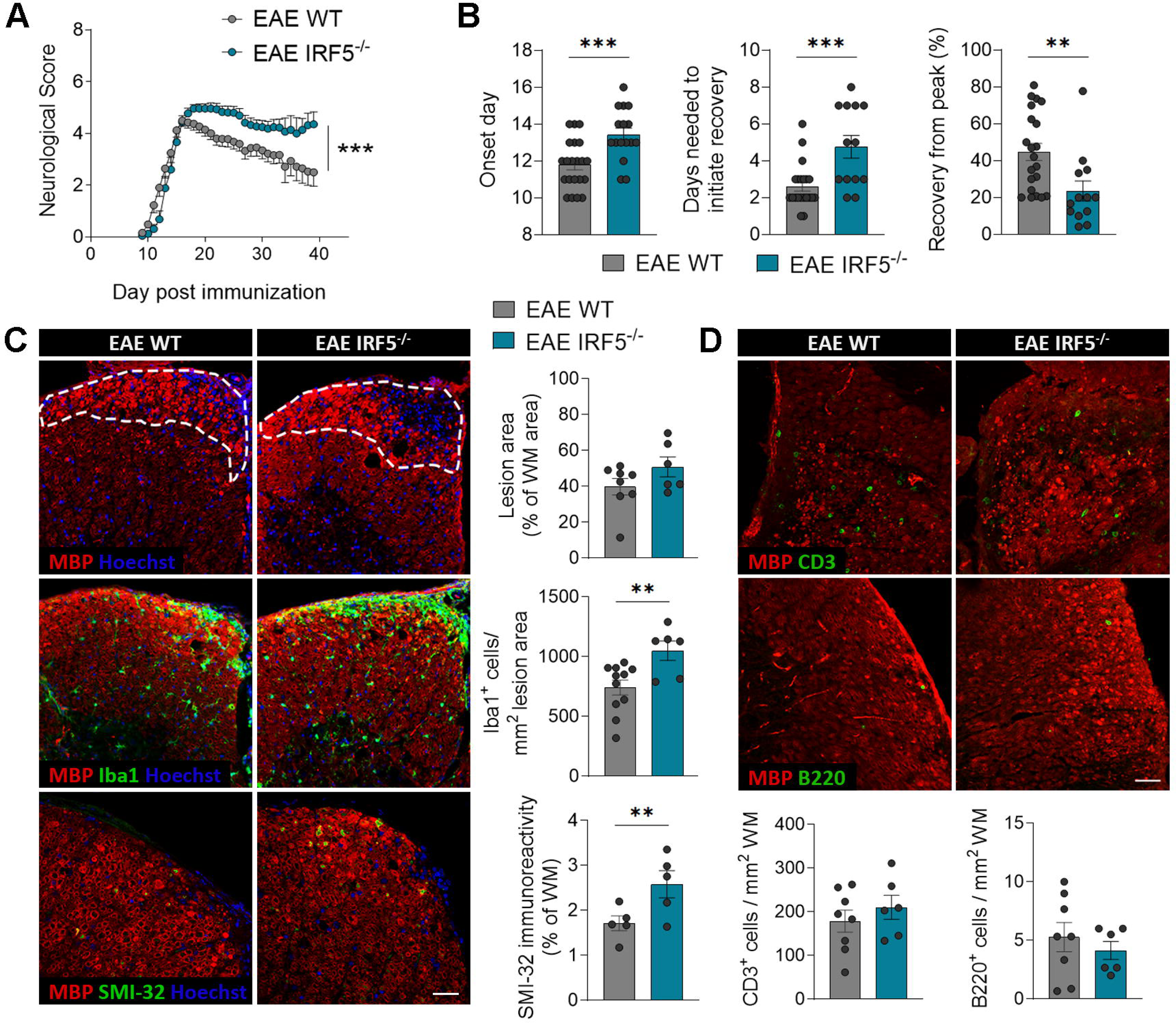
IRF5 deficiency exacerbates EAE recovery phase. **(A)** Neurological score of WT and *Irf5*^-/-^ mice (n = 10 mice; one representative experiment of three independent experiments). **(B)** Histograms showing clinical parameters associated to EAE induction (onset day) and recovery (days needed to initiate recovery and percentage of recovery from peak), in WT and *Irf5*^-/-^ mice (*n* = 13-20). **(C)** Representative images of lumbar spinal cord EAE lesions (top), Iba1 (middle) and SMI32 staining’s (bottom) in WT and *Irf5*^-/-^ mice. Immunohistochemistry was performed at 40 post-immunization. Scale bar = 30 µm. Histograms show the extent of the lesions in relation to the total white matter area of the section analyzed (*n* = 6-8) and the accumulation of Iba1^+^ microglia/macrophages (*n* = 6-11) as well axonal damage (SMI-32) in relation to the lesioned area or the total white matter area, respectively (*n* = 5). **(D)** Representative images showing the accumulation of CD3^+^ T cells and B220^+^ B cells in EAE lesions of WT and *Irf5*^-/-^ mice at day 40 post-immunization. Scale bar = 30 µm. Histograms show the number of cells normalized to the white matter area (*n* = 6-8). Data are presented as means ± SEM. **p < 0.005, ***p < 0.001.

As IRF5 is a transcription factor involved in the immune response, we checked the immune response at EAE chronic phase. We did not observe any significant difference in the accumulation of immune cells in the lesions, neither in the case of T cells nor B cells (assessed by CD3 and B220 staining, respectively; Fig. 2D) at EAE chronic phase. To delve into the immune response, we also measured the levels of CD4^+^ and CD8^+^ cells in the spinal cord, as well as the different CD4^+^ T cell subtypes in spinal cord and peripheral immune organs, by qPCR. We did not find significant alterations in the expression of *Cd4* or *Cd8* (Fig. S1A), nor in the expression of *FoxP3, Ror* and *Ifn*γ, signature genes for Treg, Th17 and Th1 cells, respectively (Fig. S1B). These results suggest that the differences observed in EAE recovery are not due to alterations in the adaptive immune response. IRF5 is known to be involved in regulating microglia/macrophage response ^12, 13^. However, the expression of different pro-inflammatory and anti-inflammatory markers of microglia, as analyzed by Fluidigm qPCR (Fig, S1C), were upregulated in *Irf5*^-/-^ mice at EAE chronic phase, but none significantly shift was detected in the pro versus anti-inflammatory profile. Altogether, these data suggest that IRF5 transcription factor is necessary to EAE recovery through a mechanism independent of its classic modulatory role of the immune response.

### IRF5 deficiency worsens LPC-induced demyelination and alters oligodendrocyte recruitment

To test whether IRF5 has a role in EAE recovery and remyelination we used a chemical-induced demyelination model, an immune-free model more relevant to study remyelination mechanisms^26^. We provoked focal demyelinating lesions in WT and *Irf5*^-/-^ mice by injecting 1% LPC into the white matter tracts of the spinal cord ^40^. Lesions were histologically analyzed after 14 days (Fig. 3A); at this timepoint, OPCs have been recruited into the lesions, have differentiated into mature oligodendrocytes and the remyelinating processes are ongoing ^26^.

**Figure 3.**
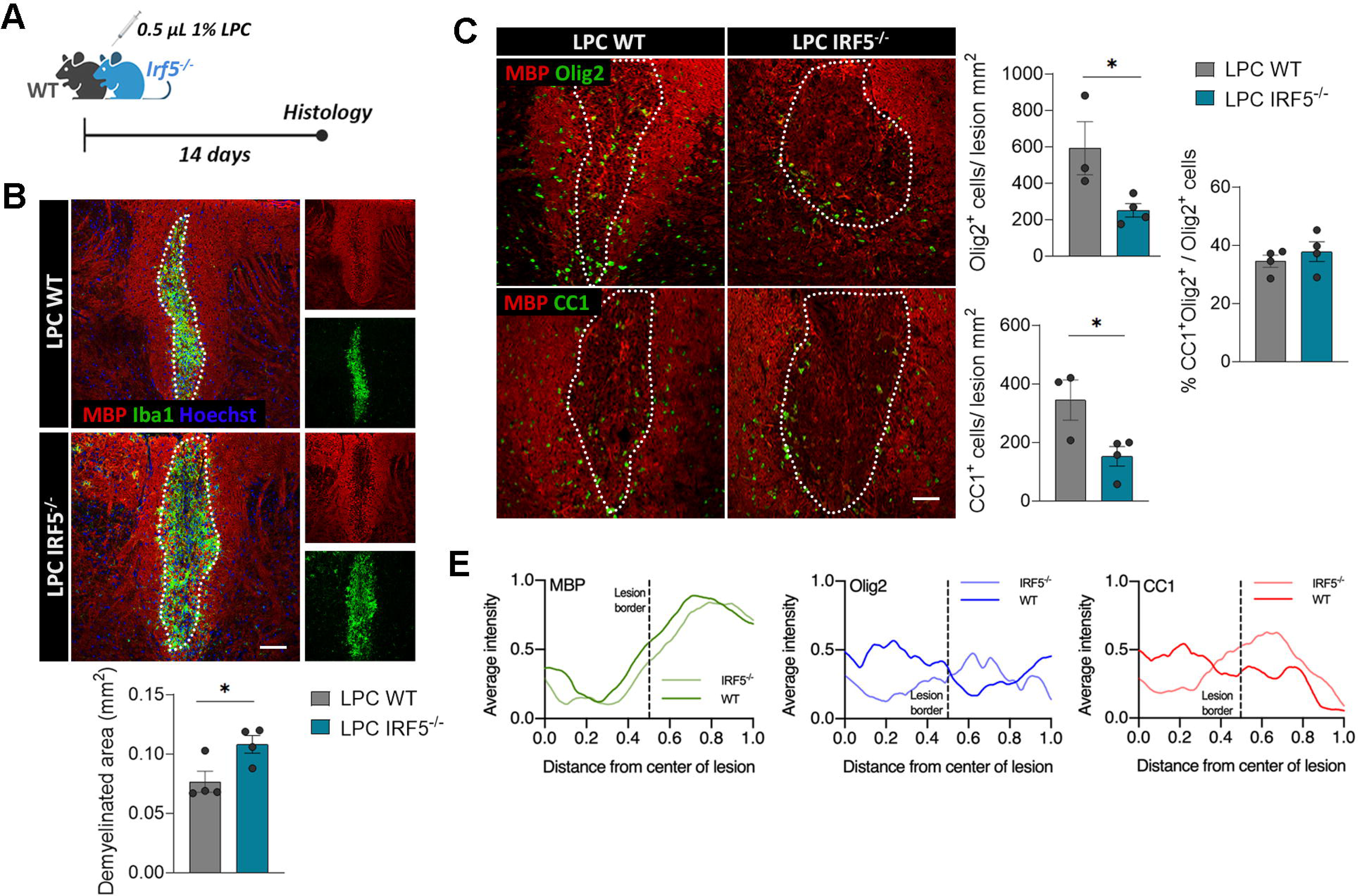
IRF5 deficiency alters remyelination after lysolecithin (LPC)-induced lesions. **(A)** Scheme showing the experimental design of LPC-induced demyelinated lesions. Analysis was performed at 14 post-injection, a time coincident with the initiation of the remyelinating response. **(B)** Representative images of MBP and Iba1 stainings in LPC lesions of WT and *Irf5*^-/-^ mice. Scale bar = 75 µm. Histogram shows the extent of demyelinated area in each mice (*n* = 4). **(C)** Assessment of the number of Olig2^+^ and CC1^+^ oligodendrocytes in LPC-induced lesions, delineated by MBP loss (*n* = 3-4). Scale bar = 75 µm. **(E)** Distribution analysis of MBP (left) and Olig2 (middle) and CC1 immunostaining, in an area comprising equal distances of lesioned and non-lesioned white matter (lesion border indicated with dotted lines; *n* = 4). Data are presented as means ± SEM. *p < 0.05.

In accordance to the EAE results, *Irf5*^-/-^ mice showed an exacerbated pathology upon LPC injections, presenting larger lesions (Fig. 3B) at 14 days post-injection. Moreover, *Irf5^-/^*^-^ mice presented more abundance of infiltrating CD3^+^ T cells in the lesions at 14 days post-injection (Fig. S2A), indicative of an aberrant or exacerbated inflammatory and immune response. In spite of these results, we did not detect more axonal damage as determined by SMI32 immunostaining (Fig. S2B). Regarding the remyelination process, we detected a decrease in the total number of oligodendrocytes, determined by the Olig2 marker, in the lesions of *Irf5*^-/-^ mice (Fig. 3C). Similarly, we found a diminished population of mature oligodendrocytes in these animals, assessed by CC1 staining (Fig. 3C); nevertheless, the proportion of myelinating CC1^+^ Olig2^+^ cells in relation to the total Olig2^+^ population was not different between genotypes (Fig. 3C). This points out to an alteration in oligodendrocyte recruitment, and not in their differentiation capacity. This feature is accompanied by an abnormal distribution of oligodendrocytes within the lesions. Both Olig2^+^ and CC1^+^ oligodendrocytes were mainly disposed in the lesion border and peri-lesion in *Irf5^-/^*^-^ mice, rather than in the lesion core delineated on the basis of MBP immunoreactivity loss (Figs. 3C and Fig. 3D).

Altogether these finding corroborate that IRF5 is necessary for a proper remyelination response, probably by regulating innate immune response such as microglia.

### Transcriptional profiling of *Irf5^-/^*^-^ microglia shows alterations in metabolism and intracellular signaling

Since microglia are essential for a proper remyelination and Irf5 controls microglia responses, we next performed bulk RNA-sequencing of FACS-sorted microglia to identify new signaling pathways regulated by *Irf5* in microglia, We FACS-sorted microglia (Cd11b^+^ CD45^low^ Ly6C^-^ population; gating strategy in Fig. 4A) from the spinal cords of control WT and *Irf5^-/^*^-^ mice (Fig. 4A). We detected a high number of differentially expressed genes between WT and *Irf5^-/^*^-^ mouse microglia (Fig. 4B; DEGs had a log (Fold Change) > 1 and adjusted p-value < 0.05).

**Figure 4.**
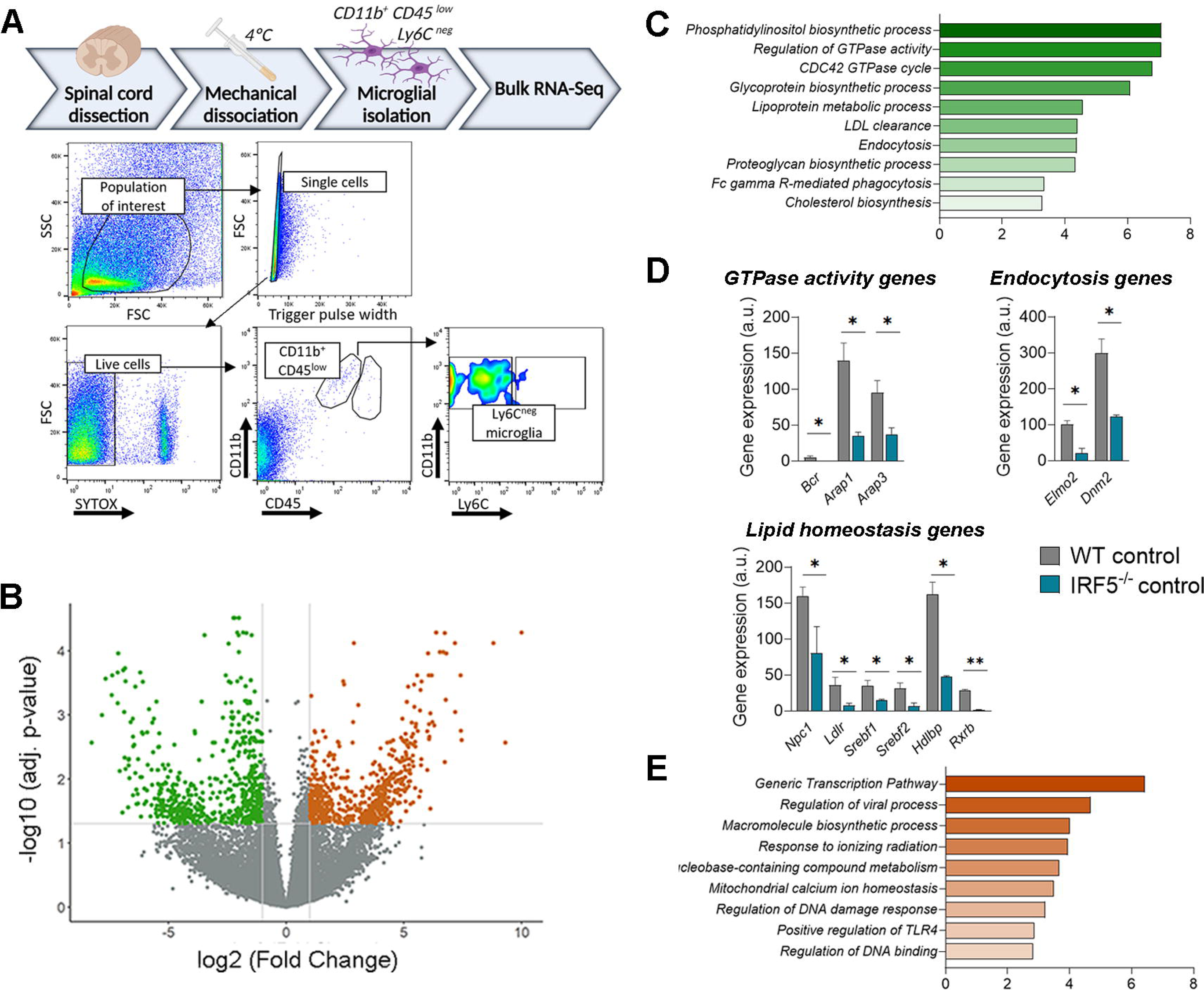
RNA sequencing of WT and *Irf5*^-/-^ microglia highlights novel roles for IRF5. **(A)** (Above) Experimental strategy for isolating spinal cord microglia at 4°C to avoid overactivation and RNA sequencing. (Below) Flow cytometry gating strategy for isolation of microglia from the spinal cord of WT and *Irf5*^-/-^ mice. **(B)** Volcano plot depicting gene expression comparison between WT and *Irf5*^-/-^ microglia. Each dot represents an individual gene. Non-significant genes are marked in gray while significant ones (log (FC) < 1 and p-value < 0.05) are marked in colour. **(C)** GO enrichment analysis of the DEGs identified between WT and *Irf5*^-/-^ microglia, showing the top GOs enriched in WT condition. Green plot shows annotations downregulated in *Irf5*^-/-^ microglia. (**D**) Histograms showing alterations in genes involved in GTPase activity (top left), endocytosis (top right) and lipid homeostasis (bottom) (*n* = 3-4). (**E**) GO enrichment analysis of the DEGs identified between WT and *Irf5*^-/-^ microglia, showing the top GOs enriched in KO condition. Orange plot shows annotations downregulated in *knock-out* microglia.

GO enrichment analysis revealed that genes downregulated in *Irf5*^-/-^ microglia associated with GTPases signaling, such as “Regulation of GTPase activity” or “CDC42 GTPase cycle” (e.g., *Bcr*, *Arap1*, *Arap3),* but mostly linked to metabolism and specifically to lipid metabolism, including “Phosphatidylinositol biosynthetic process”, “Lipoprotein metabolic process”, “LDL clearance” or “Cholesterol biosynthesis” (e.g., *Npc1*, *Ldlr, Srebf1/2*, *Hdlbp, Rxrb*…). Moreover, we also observed an association of these downregulated genes with “Endocytosis” or “Fc gamma R-mediated phagocytosis” (e.g., *Elmo2*, *Dnm2)*. All these transcriptional alterations observed in *Irf5*^-/-^ microglia could potentially be the root for the unsuccessful remyelination in mice lacking IRF5 (Fig. 4C). Predictably, other enriched GOs were related to inflammation and specific immune responses (not shown).

Conversely, IRF5-deficient microglial cells upregulated genes associated with specific immune responses such as “Regulation of viral processes” (e.g., *C3*) or “Positive regulation of TLR4” (e.g., *Wdfy1*); this is expected to be linked to the well-known role of IRF5 in immunity. Moreover, these cells also upregulated pathways associated with DNA transcription or response to DNA damage, such as “Generic Transcription Pathway”, “Response to ionizing radiation” “Regulation of DNA damage response”, “Regulation of DNA binding” or even “Macromolecule biosynthetic process” (e.g., *Bcl2*, *Brf2* or different zinc finger proteins). This outcome could be explained on the basis of the described pro-apoptotic and pro-cell cycle arrest functions of IRF5, which also participates in the p53 pathway ^41, 42^ (Fig. 4D).

The transcriptional profiling of *Irf5*^-/-^ microglia highlighted the relevance of this transcription factor in microglia; its modulatory role in intracellular pathways beyond the expected immune responses; and its potential role regulating processes required for proper remyelination.

### *Irf5*^-/-^ microglia show impaired motility in vitro but does not affect response to demyelination

GTPases in general, and Rho family of small GTPases (Rho, Rac and Cdc42) in particular, are critical regulators of the the dynamic organization of intracellular actin cytoskeleton. Thus, their associated pathways can potentially modulate a wide diversity of functions such as cellular motility or phagocytosis ^43, 44^. Indeed, Cdc42 signaling is key for both microglial migration and phagocytosis of degenerating neurons ^45^. Given the GTPase pathways downregulation observed in *Irf5*^-/-^ microglia in comparison to WT cells, we hypothesized that these functions could be compromised.

First, we performed wound healing assays on WT and *Irf5*^-/-^ microglia to test their migratory capacity and found that, after 24 hours, IRF5-deficient microglia were less efficient than WT cells repopulating the scratched area (Fig. 5A). This highlights that the altered GTPases signaling in *Irf5*^-/-^ microglia leads to abnormal motility *in vitro*, a fact that could affect microglial response to demyelination *in vivo*. However, we did not detect major changes in microglial migration towards the EAE or LPC lesions at the chronic phase (3-35 days for EAE and 14 days for LPC; see Figs 2C and 3B). To further check the impact of IRF5 deficiency in microglial migration towards demyelinating lesions, we histologically analyzed LPC-induced lesions at 4 days post-injection, a timepoint coincident with microglia/macrophage arrival (Fig. 5B) ^26^. At this stage, there was no significant difference in the migration of Iba1^+^ microglia/macrophage migration into the lesions of WT and *Irf5*^-/-^ mice. Rather, *Irf5*^-/-^ mice showed more Iba1 immunoreactivity in the demyelinated areas than WT mice did at this timepoint (Fig. 5C). Moreover, at this timepoint, there was not difference in the extent of demyelinated area between WT and *Irf5*^-/-^ mice (Fig. 5C), suggesting no differences at first demyelination responses after LPC (Fig. 5C).

**Figure 5.**
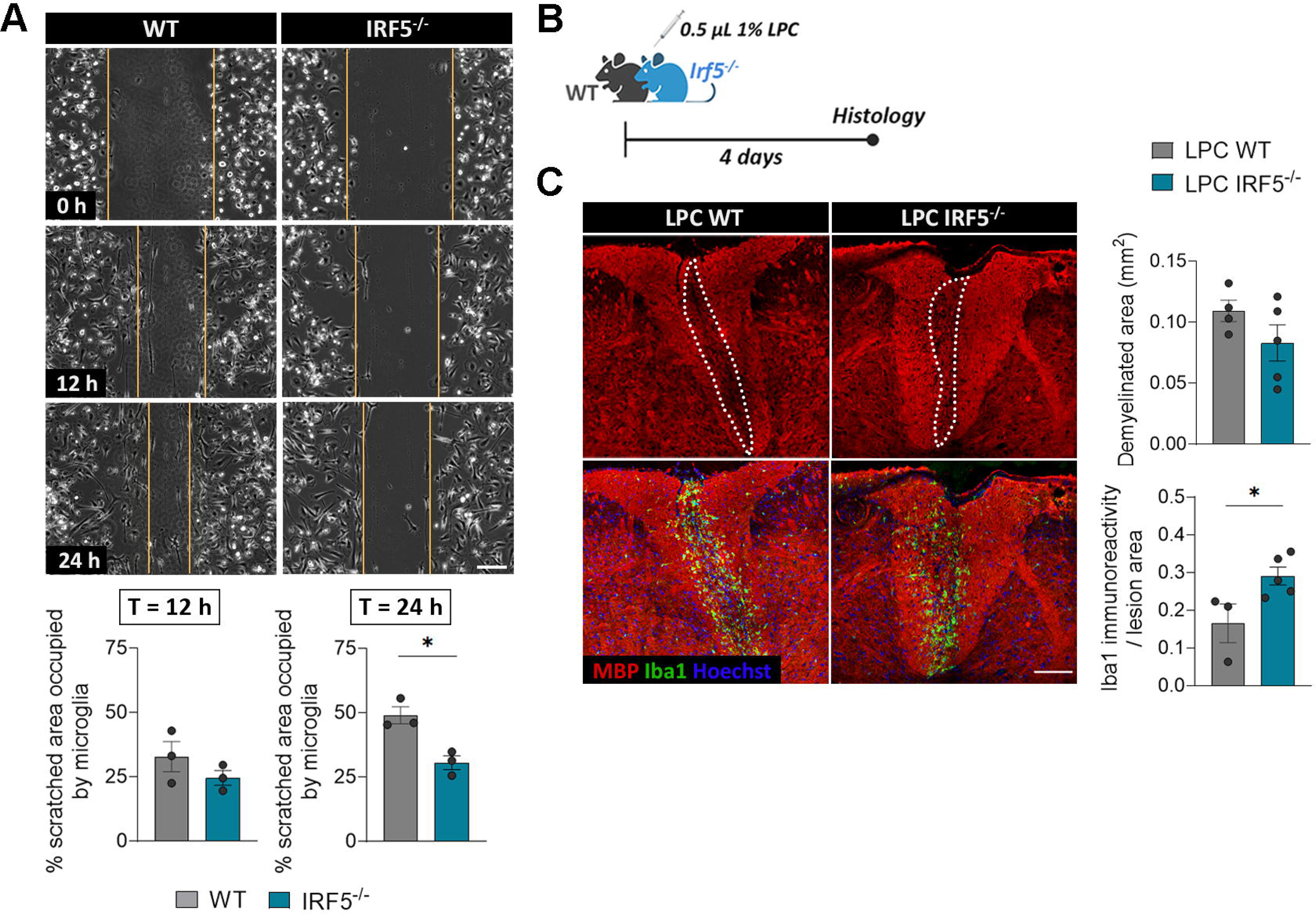
*Irf5^-/-^* microglia showed altered motility *in vitro* but not after demyelination. **(A)** Representative frames of the wound healing assay performed on WT and *Irf5*^-/-^ microglia, at the initial time of the experiment as well as after 12 and 24 hours. Yellow lines delimitate the scratched, non-occupied area at each timepoint. Scale bar = 10 µm. Histograms below show the percentage of the initially scratched area occupied by microglial cells (*n* = 3 independent experiments). **(B)** Scheme showing the experimental design for the histological analysis of microglial migration after LPC demyelinating lesions, in WT and *Irf5*^-/-^ mice, at day 4 post-injection. **(C)** Representative confocal images of LPC-induced lesions 4 days after injection, showing MBP and Iba1 immunostaining. Scale bar = 100 µm. Histograms show the extent of demyelinated area in WT and *Irf5*^-/-^ mice and Iba1^+^ immunoreactivity in relation to the lesioned area in each animal (*n* = 4-5). Data are presented as means ± SEM. *p < 0.05.

These data suggest that, although *Irf5*^-/-^ microglia showed some impairments in motility *in vitro*, this deficit does not seem to be responsible for the alterations observed in demyelinating models. Moreover, as the LPC lesions were not different between genotypes at early stages, this points out to secondary mechanisms influencing later remyelinating events.

### IRF5 deletion alters myelin clearance both *in vivo* and *in vitro*

Myelin clearance from the tissue by myeloid cells is essential for an efficient regenerative response . As we detected downregulation in the endocytic and phagocytic pathways in *Irf5* microglia by RNA sequencing, we next analyzed whether deficits in microglia phagocytosis of myelin contribute to the regeneration failure in *Irf5*^-/-^ mice.

First, we analyzed whether IRF5-deficient animals accumulated more myelin debris after demyelination. Indeed, *Irf5*^-/-^ mice showed a higher accumulation of disrupted or fragmented myelin both in EAE chronic phase and 4 days after LPC injections in the spinal cord (Fig. 6A, B). Damaged myelin yielded higher MBP immunoreactivity due to the unmasking of protein epitope ^23^. Moreover, we quantified myelin phagocytosis by microglia/macrophage in EAE lesions, assessing the presence of MBP^+^ debris inside Iba1^+^ ROIs that include the whole cytoplasm, processes and pouches using our own designed Image J macros. *Irf*5^-/-^ mice showed a higher phagocytic index (% of blobs within microglia), meaning a higher accumulation of myelin debris in microglia cells pouches or cytoplasm (Fig. 6C). We observed that myelin debris size was bigger and preferentially located in the phagocytic processes of *Irf5*^-/-^ Iba1^+^ cells. In contrast, WT microglia presented more abundance of partially degraded myelin in the cytoplasm (Fig. 6C). Thus, although *Irf5*^-/-^ microglia showed an increase in myelin phagocytosis or accumulation, the differential size and distribution of myelin debris may indicate an impairment in myelin degradation after endocytosis.

**Figure 6.**
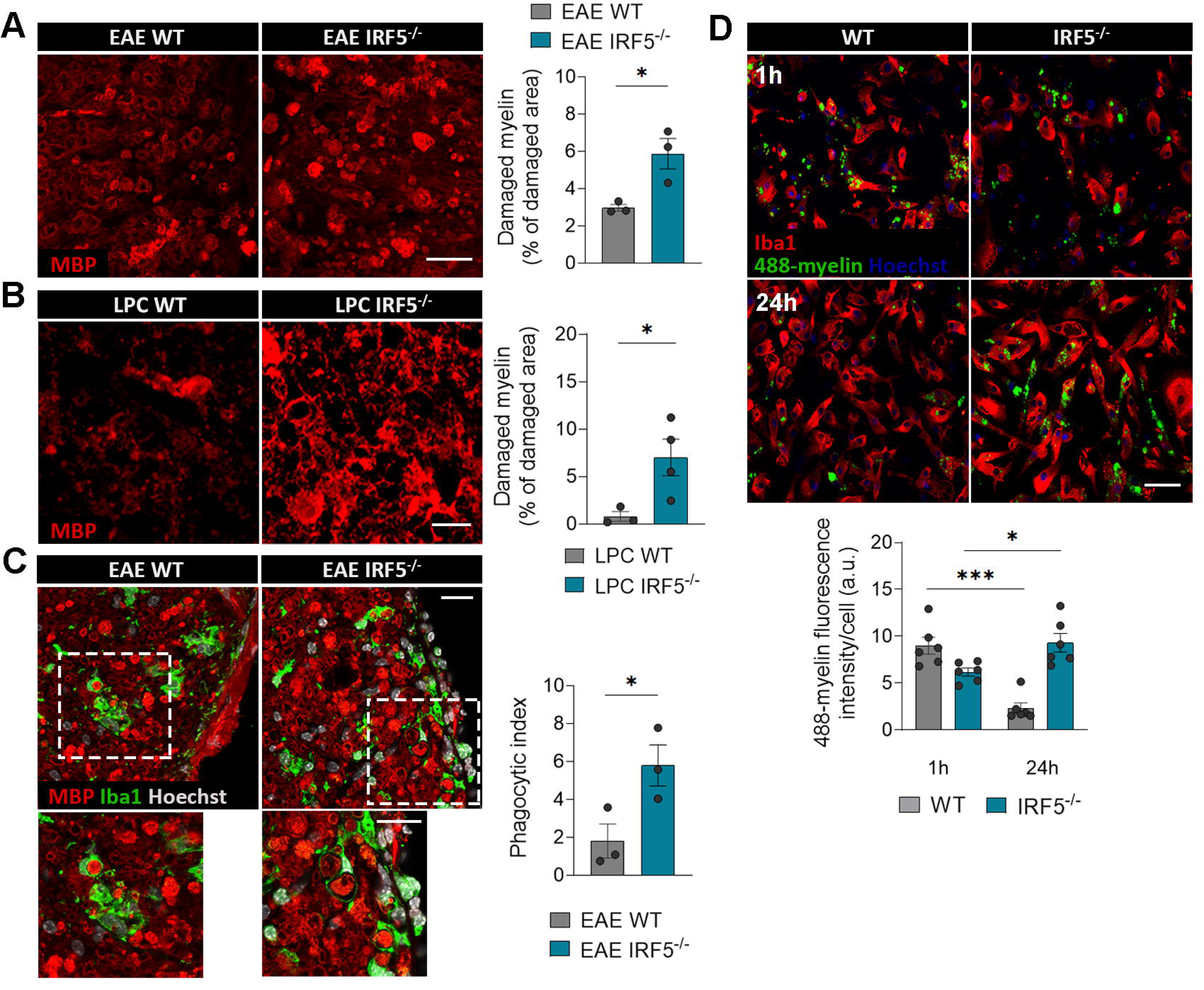
Myelin phagocytosis and degradation are altered in *Irf5^-/-^* microglia both *in vitro* and after demyelination. **(A)** Representative images of myelin debris accumulation (characterized by high MBP immunoreactivity) in WT and *Irf5*^-/-^ EAE lesions, at the recovery phase. Scale bar = 50 µm. Histogram shows the lesioned area occupied by this debris in each section (*n* = 3). **(B)** Representative images of myelin debris accumulation at day 14 post-LPC demyelinating injections, in WT and *Irf5*^-/-^ mice. Scale bar = 10 µm. Histogram shows the lesioned area occupied by this debris in each animal (*n* = 4). **(C)** Representative images of MBP and Iba1 immunostaining in spinal cord sections of WT and *Irf5*^-/-^ mice. Insets show higher magnifications of the indicated boxes. Scale bar = 20 µm. Histogram shows the phagocytic index of microglia/macrophages in these conditions (*n* = 3). **(D)** Representative images showing phagocytosis (1h) and degradation (24h) of Alexa-488 labelled-myelin by WT and *Irf5*^-/-^ microglia *in vitro*. Scale bar = 50 µm. Histogram shows the fluorescence of 488-myelin in the cells, defined as ROIs using Iba1 staining (*n* = 6 independent experiments). Data are presented as means ± SEM. *p < 0.05, ***p < 0.001.

To further explore myelin phagocytosis, myelin was isolated from adult mouse whole brain using sucrose gradient ^35^, labelled with the dye Alexa-488 and added to microglia cultures. In order to efficiently clear up myelin, microglia should internalize myelin and deliver it to lysosomes to degrade it. We monitored by confocal microscopy myelin endocytosis (1h) and myelin degradation later on (24h). We observed a significant decrease in myelin endocytosis in *Irf5*^-/-^ microglia after 1h (Fig. 6D). Moreover, while WT microglia properly degraded the internalized myelin after 24 hours, *Irf5*^-/-^ microglia showed a faulty degradatory process (Fig. 6D).

All these results suggest that IRF5 deficiency is associated to alterations in myelin phagocytosis and/or degradation, which could be the root for the regeneration failures observed in *Irf5*^-/-^ mice in response to demyelination.

### IRF5 deficiency provokes impaired lipid homeostasis and myelin metabolism after demyelination

Recently, evidence showing that not only myelin phagocytosis but also its proper intracellular processing is crucial for regeneration has arisen ^7, 8, 10^. The alterations in lipid metabolism detected in the transcriptional profile of *Irf5*^-/-^ microglia could be, thus, pivotal to explain the different responses to demyelination in animals lacking this transcription factor. In fact, genes involved in lipid endocytosis (*Ldlr*), egress of lipids from lysosomes (*Npc1*), efflux of cholesterol (*Hdlbp*), and even the transcriptional regulation of lipid homeostasis (*Srebf1* and *Srebf2*) were downregulated in *Irf5*^-/-^ microglia. These latter genes favor the generation of sterols that serve as LXR ligand and promote regenerative actions ^46^. Interestingly, the expression of the gene encoding for RXR (*Rxr*b), which dimerizes with LXR and PPAR receptors, was also downregulated in *Irf5*^-/-^ microglia (Fig. 4C). All these genes are critical modulators of lipid and cholesterol metabolism, and the deficiency of some is associated to lipid-related pathologies ^47^.These suggest that lipid homeostasis could be defective in *Irf5*^-/-^ microglia.

In order to deepen our understating in the lipid metabolism happening in *Irf5*^-/-^ microglia after demyelination, we performed MALDI-MS imaging in parallel to immunohistochemistry on the same LPC demyelinated spinal cords. After MALDI imaging, tissues were immunolabeled using antibodies to MBP and Iba1 to delineate the lesions and define the regions of interest (healthy WM, periphery and lesion core). MALDI segmentation identifies three main regions correlated with normal white matter, lesion core and periphery. The latter, higher enriched in microglia/macrophages, could help us to delineate specific changes in these population. MALDI spectra and associated lipid signatures coincident with those regions were identified and analyzed (Fig. S3A). First, we compared the lipidic profiles of healthy and lesion core areas. We observed a significant decrease in the ceramides, sulfatides, plasmalogens and phosphatydilserines (Fig. S3B), *bona fide* myelin signature lipids ^35^, in the lesion core of both WT and *Irf*5^-/-^ lesions. Myelin is characterized by a high content of these lipids ^48^, and so this decrease could be directly associated with the demyelinating processes. In addition, we also observed differences in the accumulation of phosphatidylcholines and phosphatidylethanolamines (data not shown), main components of cellular membrane, a fact that could correlate with the inflammatory events^49^. Main lipid changes detected at LPC peri-lesion represented middle values between healthy white matter and totally demyelinated white matter in the lesion core (Fig. S3C). The data suggest that the lipid signature in the periphery reggion is more probably associated to the process of demyelination than to microglia/macrophage metabolism. Interestingly, although we did not detect significant changes in the lipidic profiles between WT and *Irf5^-/-^* mice, sulfatides and phosphatidylcholines tended to show lower and higher accumulation respectively in *Irf5^-/-^* mice peri-lesion (Fig. S3D).

To more accurately analyse the impact of IRF5 in myelin metabolism in microglia, we challenged both WT and *Irf5*^-/-^ microglia with an excess of myelin (25 µg/mL) for 48 hours, and then lipids inside the cells were isolated and measured by HPLC-MS. We detected clear differences in the processing of myelin between both groups (Fig. 7A, B). Specifically, we observed significant reductions in the concentration of diverse phospholipid families including plasmalogens or phosphatidylinositols in IRF5-deficient microglia (Fig. 7B). Moreover, we detected a significant increase in the levels of all the cholesterol ester (CE) species in those cells (Fig. 7B). In contrast, no differences were observed in total free cholesterol (data not shown). A lipid ontology (LION) enrichment analysis of the changes suggested an upregulation of lipid storage in droplets, probably associated to the detected increase in CEs levels (Fig. 7C). Moreover, the LION analysis pointed out to an accumulation of lipids with high lateral diffusion that form bilayers with low thickness (Fig. 7C), suggesting increased membrane dynamics. Other enriched terms found were associated with endosomal/lysosomal lipids and glycerolipids. All these changes support a microglial impairment of intracellular lipid processing and homeostasis in the absence of IRF5, which could cause a misfunction of microglia and the detected failures in remyelination.

**Figure 7.**
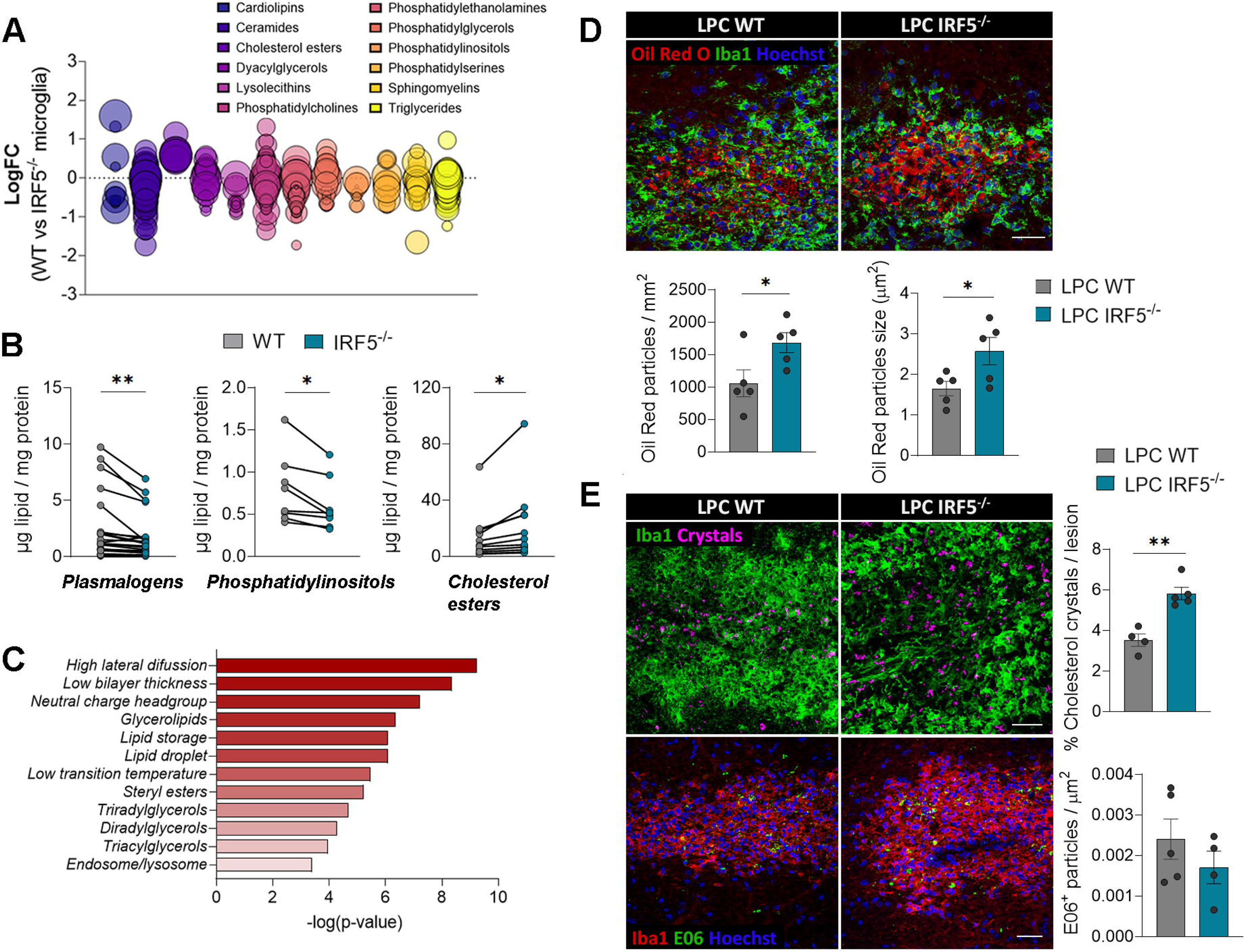
IRF5 deficiency leads to defective myelin processing and accumulation of abnormal lipid structures. **(A)** Bubble plot showing the concentration differences of specific lipids, classified by lipid classes, accumulated in WT and *Irf5*^-/-^ microglia, after 48 hour-treatment with myelin (*n* = 4 independent experiments). Each bubble represents a unique lipid species, and the size of the bubbles represents the significance of the individual lipid comparison. **(B)** Histograms showing the concentration of different plasmalogens, phosphatidylinositols and cholesterol esters in WT and *Irf5*^-/-^ microglia, after 48 hours of myelin challenge. Data are presented as means for every lipid species from 4 independent experiments. **(C)** Lipid ontology (LION) enriched terms in *Irf5*^-/-^ microglia compared to WT microglia. **(D)** Immunostaining of Oil Red O (ORO) and Iba1 in LPC-induced lesions (14 dpi) in WT and *Irf5*^-/-^ mice. Histograms show the number and size of ORO^+^ particles in the lesions (*n* = 5). Scale bar = 25 µm. **(E)** Representative images of cholesterol crystals, acquired by reflection microscopy (above), and E06^+^ (below) particles in LPC-induced lesions (14 dpi), both in WT and *Irf5*^-/-^ mice. Histograms show the percentage of lesioned area occupied by crystals and the number of EO6^+^ particles normalized to the lesion area (n = 4-5). Scale bars = 30 µm. Data are presented as means ± SEM. *p < 0.05, **p < 0.005.

HPLC-MS analysis suggest that microglia processing of myelin is altered in *Irf5*^-/-^ mice, leading to an aberrant accumulation of lipid intermediates such as CEs. Myelin-derived cholesterol is released into the extracellular space via specific transporters or it is converted to CE by the cholesterol acyltransferase and subsequently accumulated in intracellular lipid droplets (LDs). We assessed cholesterol storage after LPC-induced demyelination (14-days post-injection) using Oil Red O^+^ (ORO) staining, selective for neutral lipids. In line with the previous results, we detected an increase in the number and size of lipid bodies (LDs) in the lesions of Irf5-/- mice (Fig. 7D). As microglial defective processing of cholesterol is linked to abnormal, pathogenic formation of dense crystals ^10^, we hypothesized that our *Irf5*^-/-^ mice could also present this feature. Indeed, reflection microscopy showed an enhanced accumulation of these cholesterol crystals in the lesions of mice lacking IRF5, in comparison to WT animals (Fig. 7E). We also checked the accumulation of oxidized phosphatidylcholines (OxPCs) in LPC-demyelinated lesions. OxPCs are considered pro-degenerative factors in MS, normally being neutralized and processed by microglia ^50^. Despite the alterations in *Irf5*^-/-^ microglial myelin phagocytosis and metabolism, we did not find significant differences in the levels of OxPCs, determined by E06 staining, in WT versus *Irf5*^-/-^ lesions (Fig. 7F).

Lipid droplet accumulating microglia represent dysfunctional and proinflammatory state that could contribute to the exacerbated damage observed in *Irf5*^-/-^ after EAE. We therefore reasoned that facilitating cholesterol transport could improve EAE neurological symptoms. To test this hypothesis, we used drugs such as the GW3965 LXR agonist, that can reduce CE accumulation by upregulating the expression of ABCA1 and ABCG1 transporters ^51^, as well as 2-hydroxypropyl-β-cyclodextrin (HβCD), a molecule known to reduce intracellular cholesterol accumulation in Niemann–Pick disease type C1 disorder ^52^. We tested the impact of GW3695 and cyclodextrin in EAE pathogenesis in *Irf5*^-/-^ mice. GW3695 (20mg/kg daily; i.p. injection) and cyclodextrin (400mg/kg every 48 hours; subcutaneous injection) were administered from 10 post-immunization to avoid interfering with immune priming. GW3695 and cyclodextrin induced a significant improvement of neurological symptoms in *Irf5*^-/-^ mice (Fig. 8A). Accordingly, Oil red staining showed a significant reduction in lipid droplets accumulation into demyelinated lesions (Fig. 8B)

**Figure 8.**
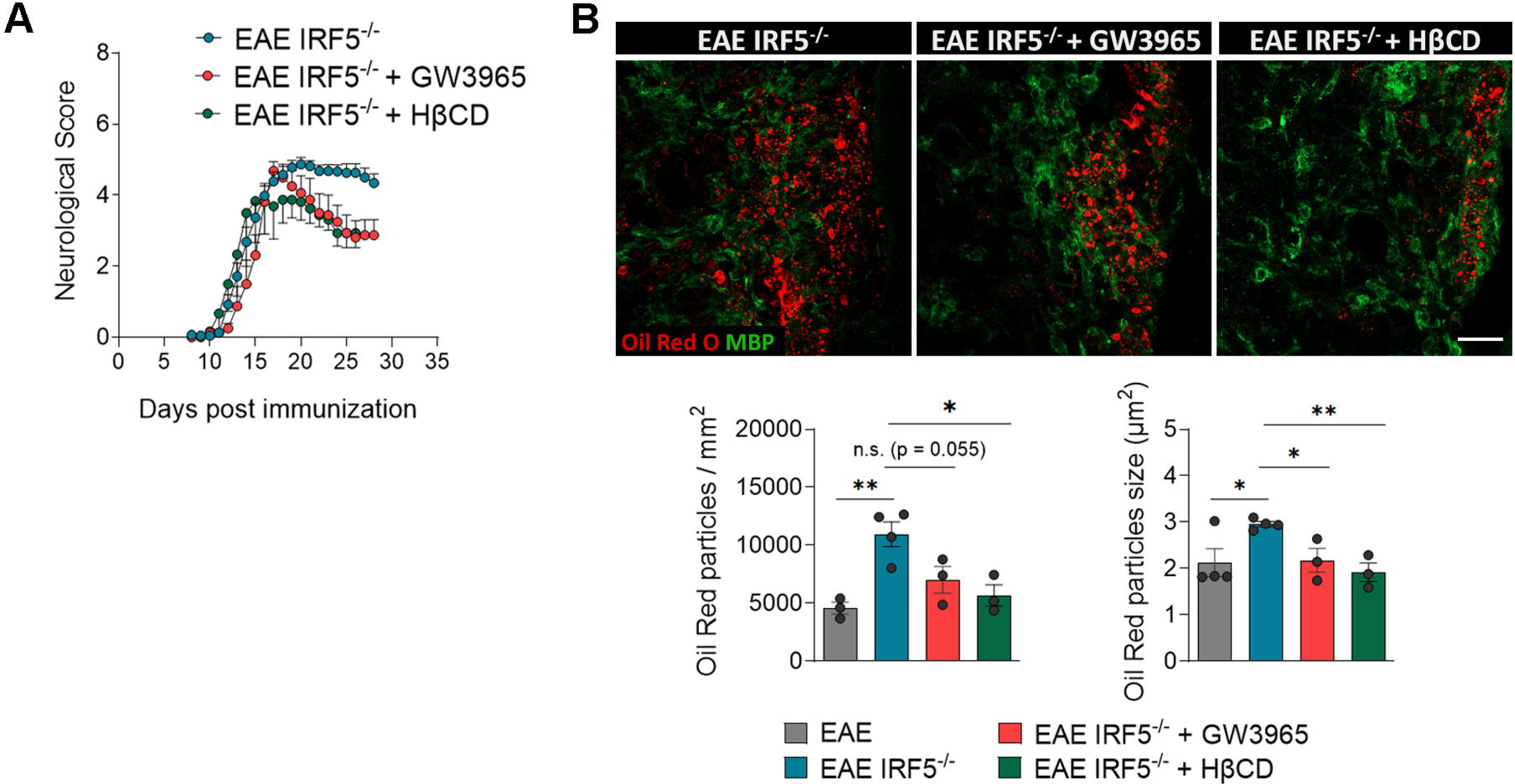
**Treatment preventing CE accumulation reverses EAE exacerbated pathology in *Irf5*^-/-^ mice**. **(A)** Neurological score of *Irf5*^-/-^ mice treated wtih saline, GW3965 (20 mg/kg; i.p.), an LXR agonist, and 2-hydroxypropyl-β-cyclodextrin (HβCD; 400 mg/kg; subcutaneous injection every 48h). Treatments started at 10 days postimmunization to avoid interfering with immune priming (n= 10). (**B**) Representative images of lipid droplets (Oil Red O staining) accumulation inside the EAE lesions in *Irf5*^-/-^ mice without treatment or treated with GW3965 and HβCD Histograms show the number and size of ORO^+^ particles in the lesions (*n* = 4-5). Scale bar = 25 µm.

Altogether these results confirm the role of IRF5 in lipid metabolism in response to demyelination; the lack of this transcription factor alters myelin processing and promotes the generation of abnormal lipid structures, hindering proper regeneration.

## DISCUSSION

Microglial lipid phagocytosis and processing after demyelination is emerging as an important focus of research, in order to promote remyelination ^7, 53, 54^. Indeed, alterations in these mechanisms have been associated to enhanced inflammatory responses and impairments in regeneration ^10, 46, 55^. Our results point out to a novel role of IRF5 modulating lipid metabolism in microglia, beside its well-described contributions orchestrating immune responses. In fact, our *Irf5^-/-^*mice showed deficient myelin processing and lipid homeostasis in response to demyelination, impeding proper remyelination, and facilitating cholesterol transport reverse these effects.

Gain of function polymorphisms in IRF5 have been associated with an increased risk of developing autoimmune diseases ^56^. IRF5 is determinant to promote disease development in a murine model of lupus erythematosus autoimmune disease ^57^. Similarly, our data showed that IRF5 play a key role in adaptive immune priming, as *Irf5*^-/-^ showed a delay in EAE onset. Our results are in accordance to previous data on literature describing the role of other IFN transcription factors in EAE development. Thus, *Irf1^-/-^, Irf3^-/-^* or *Irf8^-/-^*mice show amelioration of EAE pathogenesis ^58, 59^ or complete resistance to EAE development in the case of IRF8^39^. Unexpectedly, *Irf5*^-/-^ mice, despite this initial delay in immune priming, developed a more exacerbated EAE pathogenesis in the chronic phase. During the early phase of EAE, infiltrating monocytes and monocyte-derived macrophages as well as parenchymal microglia contribute to T cell recruitment, especially CD4^+^ T cells, into the CNS, resulting in neuronal demyelination; however, in later stages, they promote remyelination and recovery by removal of myelin debris by phagocytosis. As adaptive immune response was not altered in *Irf5*^-/-^ mice at EAE chronic phase, our data support the concept that IRF5 deletion impairs remyelination and EAE recover by a mechanism involving parenchymal microglia and/or infiltrating macrophages. Thus, IRF5 is almost selectively expressed in microglial cells in the brain and exacerbated damage and remyelination failure was also observed in the immune free model induced by LPC injections into the spinal cord. Our study points to alterations in OPCs recruitment towards LPC-induced lesions in the absence of IRF5. Moreover, we observed an increase in the number of T cells in the parenchyma of the spinal cord LPC lesions in *Irf5*^-/-^ mice, supporting the idea that myelinopathy in *Irf5*^-/-^ mice could trigger an aberrant inflammatory and immune reaction. The beneficial role of IRF5 in regeneration and remyelination is in line with studies proposing that inflammatory activation of myeloid cells is essential for this process ^11, 60^. In fact, TLR responses and even MyD88-mediated pathways, both directly linked to IRF5, are relevant for remyelination ^11^.

Our transcriptomic and lipidomic analysis point to a key role of *Irf5*^-/-^ in microglia lipid metabolism, which could explain the altered remyelination response. Normal microglial myelin clearance involve lipid metabolic pathways that eventually promote anti-inflammatory, phagocytic properties, mainly mediated by LXR/RXR activation ^8, 46^. Myelin-derived cholesterol is a key factor controlling microglia regenerative response. As cholesterol cannot be degraded, it is either release to the extracellular medium by cholesterol export transporters such as ABCA1, ABCG1 and ApoE or esterified by acyl-CoA:cholesterol acyltransferase and stored in lipid droplets, leading to foamy microglia/macrophages. *Irf5*^-/-^ microglia/macrophages showed an aberrant accumulation of lipidic structures, such as CE-containing LDs, resembling the foamy, lipid-overloaded macrophages detected in chronic MS lesions ^61^. LD-accumulating microglia represent a dysfunctional and pro-inflammatory state in the aging brain ^62^, and a similar phenotype was described in microglia lacking TREM2 ^63^. These cells abnormally enhanced LD production upon excessive myelin challenge, promoting pathogenic events such as cholesterol crystals, that block remyelination ^62–64^. Of note, recent data also suggest that LD biogenesis in phagocytes could be required for remyelination ^64^. One of the factors that could determine the beneficial versus detrimental role of LDs could be its adequate processing. Indeed, increasing lypolisis-mediated LD turnover by targeting perilipin-2, one of the main LD surface proteins, has been recently proposed as a possible theraupetic target to promote regeneration ^65^. This hypothesis is also support by the aberrant aberrant accumulation of cholesterol crystals in LDs in aged phagocytes, suggesting a faulty cholesterol metabolism that leads to maladaptive pro-inflammatory responses and failure in remyelination after LPC injections^10^. As in aged microglia, *Irf5*^-/-^ microglia/macrophages also accumulated cholesterol crystals in LPC and facilitating cholesterol efflux reversed the enhanced neurological symptoms in EAE. Our results suggest that the defective cholesterol processing in *Irf5^-/-^* mice impedes the anti-inflammatory, pro-regenerative effects of microglia/macrophages. Also, the alterations in lipid efflux could potentially dampen the capacity to recycle lipids for remyelination ^54^.

*Irf5*^-/-^ microglia also showed downregulation in genes associated with GTPases signaling, postulating these pathways as potential targets for this transcription factor. GTPases are critical modulators of actin cytoskeleton remodelling, and have been associated to microglial migratory and phagocytic capacity ^45, 66, 67^. Moreover, the loss of Rho small GTPases in microglia has been related to pro-degenerative, excitotoxic effects ^68^. Of note, the altered phosphatidylinositols (PI) metabolism detected in *Irf5*^-/-^ microglia by LC/MS can contribute to the defective GTPases signaling, given that PIs, along with other phospholipids, actively participate in these pathways in microglia, as well as in many other intracellular mechanisms ^69^. In contrast to *Irf8*^-/-^ macrophages, that show impaired migratory capacity toward the epicenter of lesions after spinal cord injury ^70^, we did not detect direct implications for the deficient GTPases signaling in *Irf5*^-/-^ macrophages’ motility after demyelination. Nevertheless, the hijacked phagocytic capacity in response to myelin debris can be secondary to the alterations in this pathway, and not only due to the intracellular lipid overdoses.

The findings in this work point to a novel role of IRF5 favouring microglial remyelinating events, through the modulation of lipid metabolism and phagocytosis. On one hand, its deficiency is potentially beneficial at early demyelination. Nevertheless, as its expression in microglia is downregulated during the pathology, enhancing microglial IRF5 activity could represent a potential therapeutic target to promote proper myelin clearance and pro-regenerative events at late stages.

## ACKNOWLEDGMENTS

We thank Prof. Tak W. Mak from the Princess Margaret Cancer Centre, UHN (Toronto, Canada) for providing us with *Irf5*^-/-^ mice. We kindly acknowledge the SGIKER Facilities at the University of the Basque Country and facilities at Achucarro Basque Center for Neuroscience for the technical support. This work was supported by the Spanish Ministry of Economy and Competitiveness, (SAF2016-75292-R), Spanish Ministry of Science and Innovation Competitiveness (PID2019-109724RB-I00 and MCIN/AEI/10.13039/501100011033), ARSEP Foundation, Centro de Investigación Biomédica en Red, Enfermedades Neurodegenerativas (CIBERNED) (grant no. CB06/05/0076) and ERDF “A way to make Europe” (RTI2018-099267-B-I00). AM, AZ, GPM, and IT and received predoctoral fellowships from the Spanish Ministry of Education and Science (AM), the Basque Goverment (GPM) and the University of the Basque Country EHU/UPV (AZ), respectively.

## AUTHOR CONTRIBUTIONS

A.M. and M.D. contributed to the conception and design of the study, data acquisition and analysis and data interpretation. M. D. supervised the study. A.Z., G.P.M., I.T and F.S. contributed to data acquisition and analysis. I.C., O.F. and J.A.F. contributed to lipid imaging and analysis. M.K., S.K. and B.E. contributed to RNAseq analysis. A.S. performed phagocityc analysis in vivo. V.T. performed lysolecithin-induced lesions.C.M. contributed to EAE experiments. A.M. and M.D. wrote the paper with input from most authors. All authors have approved the submitted version.

---

**Figure S1.**
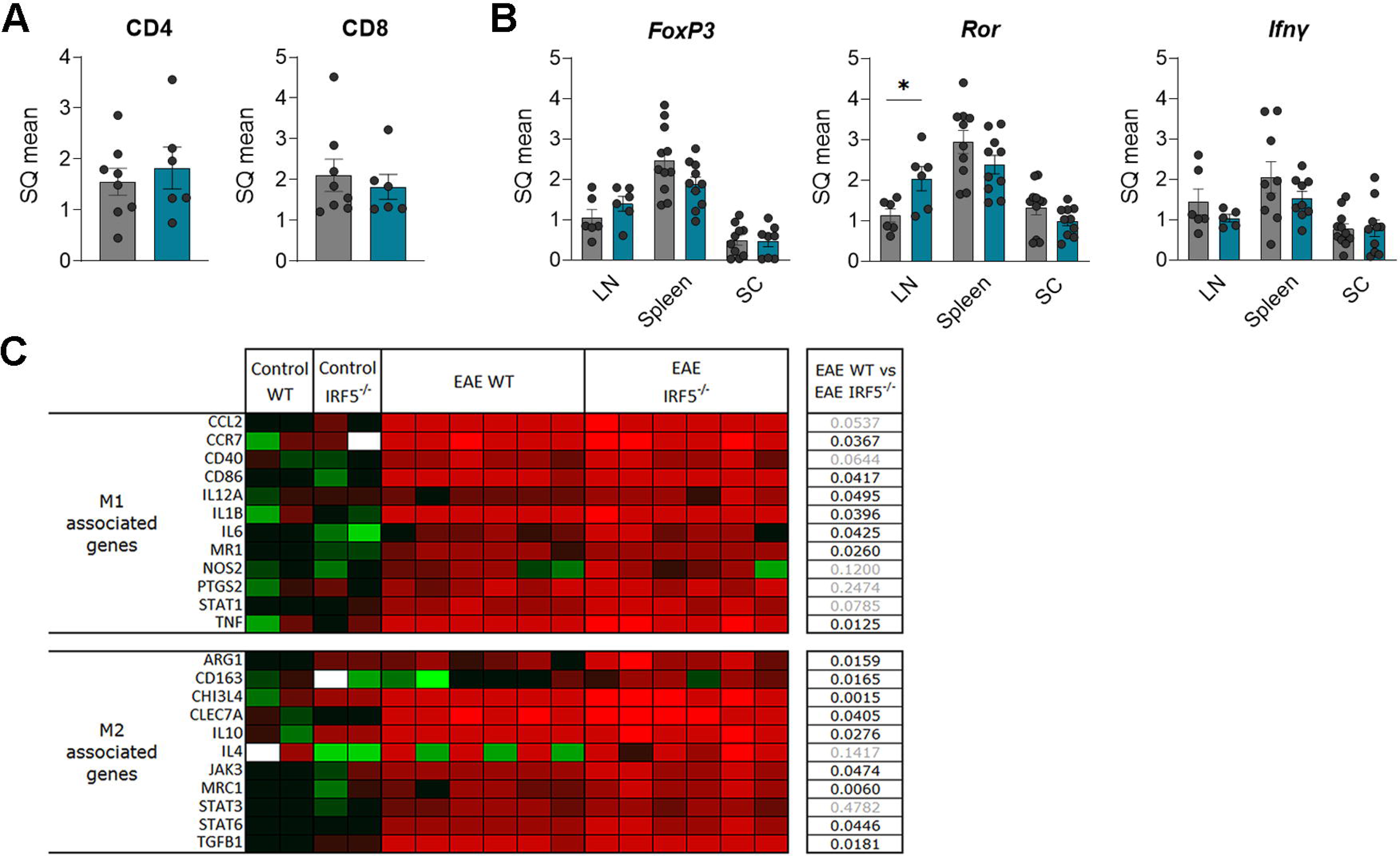
IRF5 deficiency has no impact on immune response. **(A)** Relative mRNA expression of *Cd4* and *Cd8* in the spinal cord of WT and *Irf5*^-/-^ mice at EAE chronic phase (*n* = 6-8). (**B**) Relative mRNA expression of *Foxp3* (Tregs), *Ifn*γ (Th1) and *Ror* (Th17) in lymph nodes (LN), spleen and spinal cord of WT and *Irf5*^-/-^ mice at EAE chronic phase (*n* = 6-8). (**C**) Heatmap showing significant changes in pro-inflammatory and anti-inflammatory mRNA expression in total RNA isolated from the spinal cord of WT and *Irf5*^-/-^ mice at EAE chronic phase (*n* = 6). Tables indicate statistical significance between EAE WT and EAE *Irf5*^-/-^ mice. Data were analyzed by unpaired Student’s t-test. * p<0.05.

**Figure S2.**
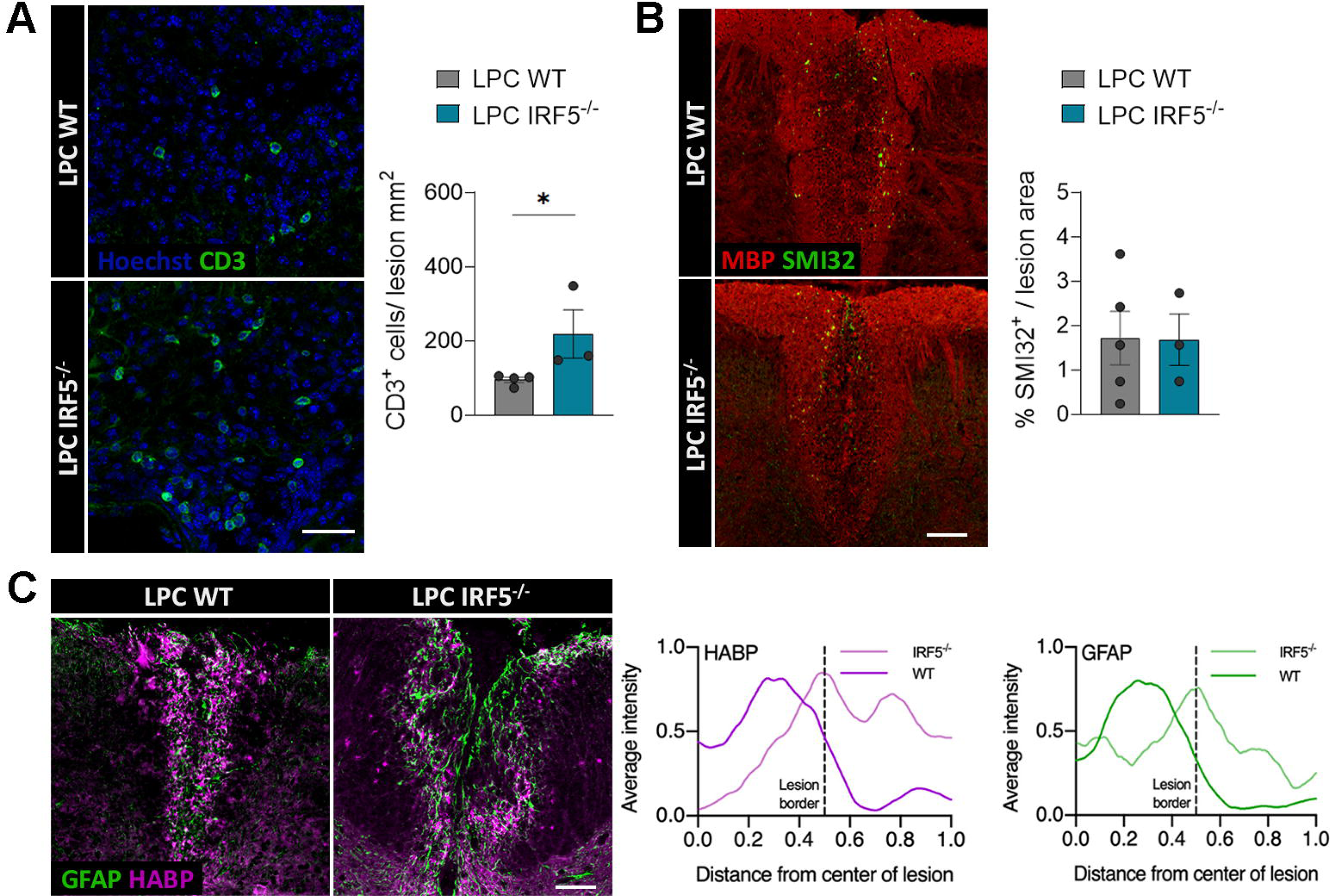
IRF5 deficiency alters T cells response after lysolecithin (LPC)-induced demyelination. (**A, B**) Spinal cord sections of wild type (WT; n = 5) and *Irf5*^-/-^ (n = 6) mice 14 days after LPC injection immunolabelled for CD3 T cell marker (**A**) and for SMI32 and MBP (**B**), markers of axonal damage and myelin. Scale bars = 30 and 75 µm, respectively. Histograms shows means ± SEM (n = 3-5). Statistic was performed with Student’ t-test. \**p* <0.05.

**Figure S3.**
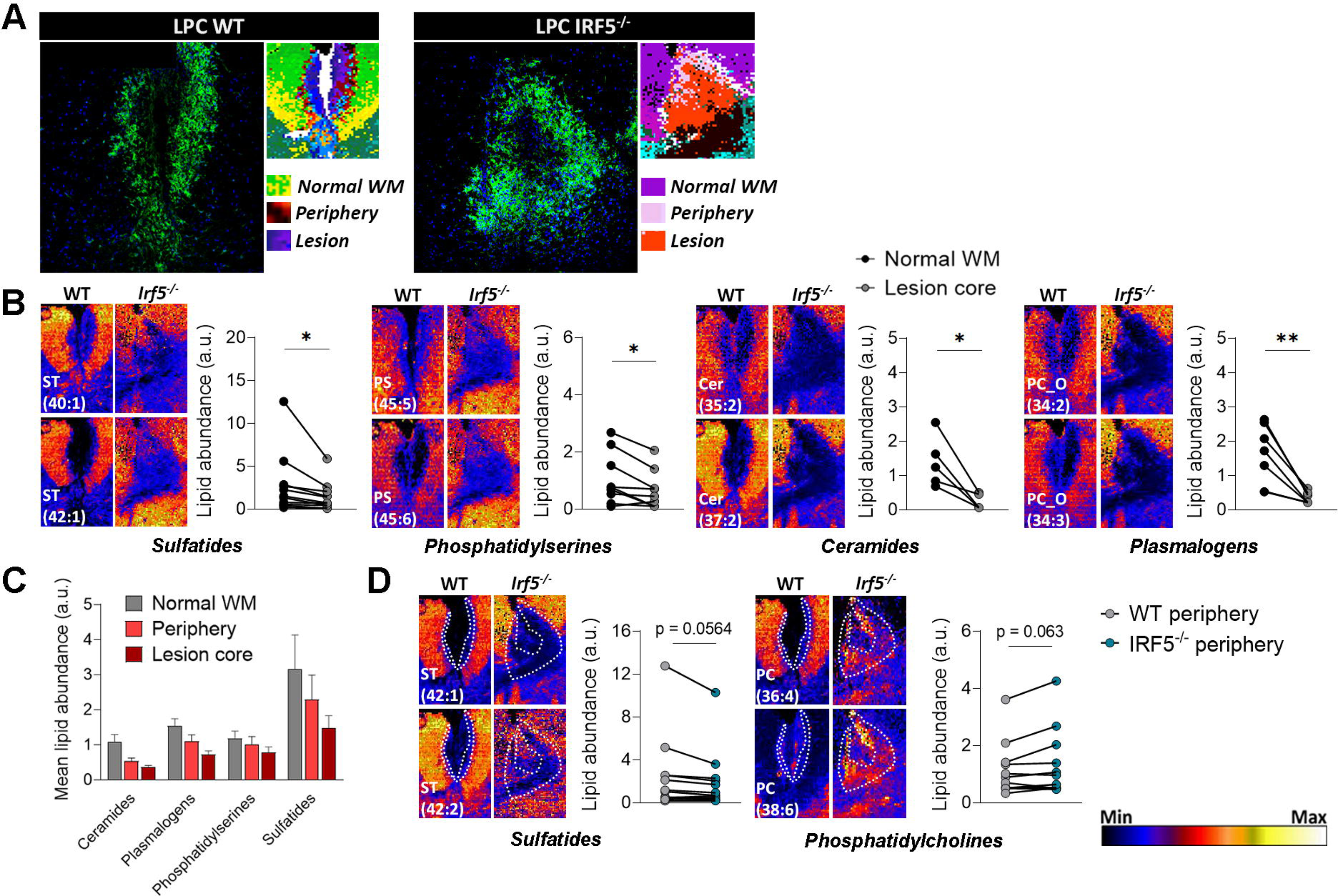
MALDI-IMS profiling of lipids in lysolecithin (LPC)-induced demyelination. (A) (**A**) Representative immunohistochemical images of LPC lesions (14 dpi), identified on the basis of Iba1 accumulation as well as MBP loss (not shown), and parallel lipid segmentation obtained of these lesions, following k-means clustering method. Of note, there are different lipid profiles corresponding to the lesion core, the periphery and the normal WM. (**B**) Representative MALDI mass spectrometry imaging (MSI) maps of major sulfatides (ST; 40:1 and 42:1), phosphatidylserines (PS; 45:5 and 45:6), ceramides (Cer; 35:2 and 37:2) and plasmalogens (PC_O; 34:2 and 34:3) in the WT and *Irf5*^-/-^ lesions. Histograms shows the difference of abundance of different families of lipids in the normal WM and the lesion core. Each point in the graphs represent a single lipid species analyzed by MALDI. (**C**) Heatmaps showing the mean lipid concentration of the different ST, PS, Cer and plasmalogens identified in the normal WM, the periphery and the lesion core. (**D**) Representative MALDI mass spectrometry imaging (MSI) maps of major sulfatides (ST; 42:1 and 42:2) and phosphatidylcholines (PC; 36:4 and 38:6) in the WT and *Irf5*^-/-^ lesions. Histograms shows the difference of abundance of different families of lipids in the WT and *Irf5*^-/-^ lesion periphery. Lipid abundance scale for MALDI-MSI images is present in the bottom right corner of the figure. Data are presented as means for every lipid species from n = 5 different mice. *p < 0.05, **p < 0.005.

## REFERENCES

1. Dendrou, C. A., Fugger, L. & Friese, M. A. Immunopathology of multiple sclerosis. Nat. Rev. Immunol. 15, 545–558 (2015).

2. Filippi, M. et al. Multiple sclerosis. Nat. Rev. Dis. Prim. 1–27 (2018).

3. Franklin, R. J. M. & Ffrench-Constant, C. Regenerating CNS myelin - From mechanisms to experimental medicines. Nat. Rev. Neurosci. 18, 753–769 (2017).

4. Neumann, H., Kotter, M. R. & Franklin, R. J. M. Debris clearance by microglia: An essential link between degeneration and regeneration. Brain 132, 288–295 (2009).

5. Plemel, J. R., Manesh, S. B., Sparling, J. S. & Tetzlaff, W. Myelin inhibits oligodendroglial maturation and regulates oligodendrocytic transcription factor expression. Glia 61, 1471–1487 (2013).

6. Kotter, M. R., Li, W. W., Zhao, C. & Franklin, R. J. M. Myelin impairs CNS remyelination by inhibiting oligodendrocyte precursor cell differentiation. J. Neurosci. 26, 328–332 (2006).

7. Franklin, R. J. M. & Simons, M. CNS remyelination and inflammation: From basic mechanisms to therapeutic opportunities. Neuron 1–17 (2022) doi:10.1016/j.neuron.2022.09.023.

8. Bogie, J. F. J. et al. Myelin-Derived Lipids Modulate Macrophage Activity by Liver X Receptor Activation. PLoS One 7, 1–10 (2012).

9. Berghoff, S. A. et al. Microglia facilitate repair of demyelinated lesions via post-squalene sterol synthesis. Nat. Neurosci. 24, 47–60 (2021).

10. Cantuti-Castelvetri, L. et al. Defective cholesterol clearance limits remyelination in the aged central nervous system. Science 359, 684–688 (2018).

11. Cunha, M. I. et al. Pro-inflammatory activation following demyelination is required for myelin clearance and oligodendrogenesis. J. Exp. Med. 217, (2020).

12. Tamura, T., Yanai, H., Savitsky, D. & Taniguchi, T. The IRF family transcription factors in immunity and oncogenesis. Annu. Rev. Immunol. 26, 535–584 (2008).

13. Takaoka, A. et al. Integral role of IRF-5 in the gene induction programme activated by Toll-like receptors. Nature 434, 243–249 (2005).

14. Krausgruber, T. et al. IRF5 promotes inflammatory macrophage polarization and T H1-TH17 responses. Nat. Immunol. 12, 231–238 (2011).

15. Saliba, D. G. et al. IRF5:RelA interaction targets inflammatory genes in macrophages. Cell Rep. 8, 1308–1317 (2014).

16. Almuttaqi, H. & Udalova, I. A. Advances and challenges in targeting IRF5, a key regulator of inflammation. FEBS J. 286, 1624–1637 (2019).

17. Masuda, T. et al. Transcription factor IRF5 drives P2X4R+-reactive microglia gating neuropathic pain. Nat. Commun. 5, 1–11 (2014).

18. Sigurdsson, S. et al. 1 Yamawaki M, Kitagaki S, Anada M, Hashimoto S. Enantioselective intramolecular C-H amidation of sulfamate esters catalyzed by chiral dirhodium(II) carboxylates. Heterocycles 2006; 69: 527–537. Am. J. Hum. Genet. 76, 528–537 (2005).

19. Shimane, K. et al. A single nucleotide polymorphism in the IRF5 promoter region is associated with susceptibility to rheumatoid arthritis in the Japanese population. Ann. Rheum. Dis. 68, 377–383 (2009).

20. Kristjansdottir, G. et al. Interferon regulatory factor 5 (IRF5) gene variants are associated with multiple sclerosis in three distinct populations. J. Med. Genet. 45, 362–369 (2008).

21. Vandenbroeck, K. et al. Validation of IRF5 as multiple sclerosis risk gene: Putative role in interferon beta therapy and human herpes virus-6 infection. Genes Immun. 12, 40–45 (2011).

22. Masuda, T. et al. Transcription factor IRF5 drives P2X4R+-reactive microglia gating neuropathic pain. Nat. Commun. 5, 1–11 (2014).

23. Zabala, A., et al. P2X4 receptor controls microglia activation and favors remyelination in autoimmune encephalitis. EMBO Mol. Med. 10, e8743 (2018).

24. Vallejo-Illarramendi, A., Domercq, M., Pérez-Cerdá, F., Ravid, R. & Matute, C. Increased expression and function of glutamate transporters in multiple sclerosis. Neurobiol. Dis. 21, 154–164 (2006).

25. Matute, C. et al. P2X7 receptor blockade prevents ATP excitotoxicity in oligodendrocytes and ameliorates experimental autoimmune encephalomyelitis. J. Neurosci. 27, 9525–9533 (2007).

26. Tepavčevic☐, V., et al. Early netrin-1 expression impairs central nervous system remyelination. Ann. Neurol. 76, 252–268 (2014).

27. Aigrot, M. et al. Genetically modified macrophages accelerate myelin repair. EMBO Mol. Med. 14, (2022).

28. Vandesompele, J. et al. Accurate normalization of real-time quantitative RT-PCR data by geometric averaging of multiple internal control genes. Genome Biol. 3, RESEARCH0034 (2002).

29. Robinson, M. D., McCarthy, D. J. & Smyth, G. K. edgeR: A Bioconductor package for differential expression analysis of digital gene expression data. Bioinformatics 26, 139–140 (2009).

30. Huang, D. W., Sherman, B. T. & Lempicki, R. A. Systematic and integrative analysis of large gene lists using DAVID bioinformatics resources. Nat. Protoc. 4, 44–57 (2009).

31. Zhou, Y. et al. Metascape provides a biologist-oriented resource for the analysis of systems-level datasets. Nat. Commun. 10, (2019).

32. Thomas, A., Charbonneau, J. L., Fournaise, E. & Chaurand, P. Sublimation of new matrix candidates for high spatial resolution imaging mass spectrometry of lipids: Enhanced information in both positive and negative polarities after 1,5-diaminonapthalene deposition. Anal. Chem. 84, 2048–2054 (2012).

33. Domercq, M. et al. System xc- and Glutamate Transporter Inhibition Mediates Microglial Toxicity to Oligodendrocytes. J. Immunol. 178, 6549–6556 (2007).

34. Bezzi, P. et al. CXCR4-activated astrocyte glutamate release via TNFa: Amplification by microglia triggers neurotoxicity. Nat. Neurosci. (2001) doi:10.1038/89490.

35. Norton, W. T. & Poduslo, S. Isolation and Characterization of Myelin. J. Neurochem. 21, 147–195 (1973).

36. Bligh, E. . & Dyer, W. . A rapid method of total lipid extraction and purification. Can. J. Biochem. Physiol. 37, 911–917 (1959).

37. Molenaar, M. R. et al. LION/web: A web-based ontology enrichment tool for lipidomic data analysis. Gigascience 8, 1–10 (2019).

38. Masuda, T. et al. Spatial and temporal heterogeneity of mouse and human microglia at single-cell resolution. Nature 566, 388–392 (2019).

39. Yoshida, Y. et al. The Transcription Factor IRF8 Activates Integrin-Mediated TGF-β Signaling and Promotes Neuroinflammation. Immunity 40, 187–198 (2014).

40. Jeffery, N. D. & Blakemore, W. F. Remyelination of mouse spinal cord axons demyelinated by local injection of lysolecithin. J. Neurocytol. 24, 775–781 (1995).

41. Barnes, B. J., Kellum, M. J., Pinder, K. E., Frisancho, J. A. & Pitha, P. M. Interferon regulatory factor 5, a novel mediator of cell cycle arrest and cell death. Cancer Res. 63, 6424–6431 (2003).

42. Hu, G., Mancl, M. E. & Barnes, B. J. Signaling through IFN regulatory factor-5 sensitizes p53-deficient tumors to DNA damage-induced apoptosis and cell death. Cancer Res. 65, 7403–7412 (2005).

43. Hanna, S. & El-Sibai, M. Signaling networks of Rho GTPases in cell motility. Cell. Signal. 25, 1955–1961 (2013).

44. Mao, Y. & Finnemann, S. C. Regulation of phagocytosis by Rho GTPases. Small GTPases 6, 89–99 (2015).

45. Barcia, C. et al. ROCK/Cdc42-mediated microglial motility and gliapse formation lead to phagocytosis of degenerating dopaminergic neurons in vivo. Sci. Rep. 2, 1–13 (2012).

46. Berghoff, S. A. et al. Microglia facilitate repair of demyelinated lesions via post-squalene sterol synthesis. Nat. Neurosci. 24, 47–60 (2021).

47. Luo, J., Yang, H. & Song, B. L. Mechanisms and regulation of cholesterol homeostasis. Nat. Rev. Mol. Cell Biol. 21, 225–245 (2020).

48. Aggarwal, S., Yurlova, L. & Simons, M. Central nervous system myelin: Structure, synthesis and assembly. Trends Cell Biol. 21, 585–593 (2011).

49. Blank, M., Enzlein, T. & Hopf, C. LPS-induced lipid alterations in microglia revealed by MALDI mass spectrometry-based cell fingerprinting in neuroinflammation studies. Sci. Rep. 12, 1–13 (2022).

50. Dong, Y. et al. Oxidized phosphatidylcholines found in multiple sclerosis lesions mediate neurodegeneration and are neutralized by microglia. Nat. Neurosci. 24, 489–503 (2021).

51. Yang, T.-M. et al. Targeting macrophages in atherosclerosis using nanocarriers loaded with liver X receptor agonists: A narrow review. Front. Mol. Biosci. 10, 1–16 (2023).

52. Ottinger, E. et al. Collaborative Development of 2-Hydroxypropyl-&#946;-Cyclodextrin for the Treatment of Niemann-Pick Type C1 Disease. Curr. Top. Med. Chem. 14, 330–339 (2014).

53. Bogie, J. F. J., Stinissen, P. & Hendriks, J. J. A. Macrophage subsets and microglia in multiple sclerosis. Acta Neuropathol. 128, 191–213 (2014).

54. Berghoff, S. A., Spieth, L. & Saher, G. Local cholesterol metabolism orchestrates remyelination. Trends Neurosci. 45, 272–283 (2022).

55. Tepavčević, V. & Lubetzki, C. Oligodendrocyte progenitor cell recruitment and remyelination in multiple sclerosis: the more, the merrier? Brain 145, 4178–4192 (2022).

56. Eames, H. L., Corbin, A. L. & Udalova, I. A. Interferon regulatory factor 5 in human autoimmunity and murine models of autoimmune disease. Transl. Res. 167, 167–182 (2016).

57. Pellerin, A. et al. Monoallelic IRF5 deficiency in B cells prevents murine lupus. JCI Insight 6, 1–18 (2021).

58. Tada, B. Y., Ho, A., Matsuyama, T. & Mak, T. W. Reduced Incidence and Severity of Antigen-induced Regulatory Factor-1. J Exp Med 185, 231–238 (1997).

59. Fitzgerald, D. C. et al. Interferon regulatory factor (IRF) 3 is critical for the development of experimental autoimmune encephalomyelitis. J. Neuroinflammation 11, 1–7 (2014).

60. Bosch-Queralt, M. et al. Diet-dependent regulation of TGFβ impairs reparative innate immune responses after demyelination. Nat. Metab. 3, 211–227 (2021).

61. Grajchen, E., Hendriks, J. J. A. & Bogie, J. F. J. The physiology of foamy phagocytes in multiple sclerosis. Acta Neuropathol. Commun. 6, 124 (2018).

62. Marschallinger, J. et al. Lipid-droplet-accumulating microglia represent a dysfunctional and proinflammatory state in the aging brain. Nat. Neurosci. 23, 194–208 (2020).

63. Nugent, A. A. et al. TREM2 Regulates Microglial Cholesterol Metabolism upon Chronic Phagocytic Challenge. Neuron 105, 837–854.e9 (2020).

64. Gouna, G. et al. TREM2-dependent lipid droplet biogenesis in phagocytes is required for remyelination. J. Exp. Med. 218, (2021).

65. Loix, M. et al. Perilipin-2 limits remyelination by preventing lipid droplet degradation. Cell. Mol. life Sci. 79, 515 (2022).

66. Rong, Z. et al. Activation of FAK/Rac1/Cdc42-GTPase signaling ameliorates impaired microglial migration response to Aβ42 in triggering receptor expressed on myeloid cells 2 loss-of-function murine models. FASEB J. 34, 10984–10997 (2020).

67. Scheiblich, H. & Bicker, G. Regulation of Microglial Phagocytosis by RhoA/ROCK-Inhibiting Drugs. Cell. Mol. Neurobiol. 37, 461–473 (2017).

68. Socodato, R. et al. Microglia Dysfunction Caused by the Loss of Rhoa Disrupts Neuronal Physiology and Leads to Neurodegeneration. Cell Rep. 31, (2020).

69. Tokizane, K. et al. Phospholipid localization implies microglial morphology and function via Cdc42 in vitro. Glia 65, 740–755 (2017).

70. Kobayakawa, K. et al. Macrophage centripetal migration drives spontaneous healing process after spinal cord injury. Sci. Adv. 5, (2019).

